# Treg-derived IκBζ promotes their conversion into Th2-like effectors and drives type 2 inflammation via BATF

**DOI:** 10.64898/2026.02.23.707402

**Authors:** Tanja Kübelbeck, Ana-Marija Kulis-Mandic, Antonia Kolb, Emmanouil Stylianakis, Zeynep Ergün, Delia Mihaela Mihoc, Sara Salome Helbich, Kartikeya Singh, Matthias Klein, Niklas Beumer, Berenice Fischer, Stephan Hailfinger, Klaus Schulze-Osthoff, Ari Waisman, Michael Delacher, Vigo Heissmeyer, Nadine Hoevelmeyer, Sebastian Reuter, Daniela Kramer

**Author notes:** corresponding authors to whom correspondence should be addressed: Prof. Dr. Daniela Kramer, University Medical Center of the Johannes-Gutenberg University of Mainz, Department of Dermatology, Langenbeckstr. 1, 55131 Mainz, Germany, Tel.: +496131/17-5731. Shared first authorship (Tanja Kübelbeck, Ana-Marija Kulis-Mandic). Shared last authorship (Sebastian Reuter, Daniela Kramer). M.D. received personal fees from Odyssey Therapeutics outside the submitted work. The other authors declare no competing interests.

## Abstract

Regulatory T cells (Treg cells) maintain peripheral immune tolerance but display considerable plasticity in peripheral tissues. The molecular mechanisms governing their function and plasticity, particularly under inflammatory conditions, remain poorly defined. Here, we identify the NF-κB-associated transcriptional cofactor IκBζ as a critical regulator of Treg cell plasticity and function. Enforced expression of IκBζ in Treg cells triggered the excessive expansion of functionally impaired Treg cells, resulting in lymphadenopathy, splenomegaly, and systemic type 2 inflammation, most prominently in the lung. Mechanistically, IκBζ modified BATF expression and function, thereby driving the cell-intrinsic production of Th2-associated cytokines by Treg cells. Conversely, Treg-specific deletion of IκBζ constrained IL-33-mediated expansion of tissue Treg cells and surprisingly attenuated type 2 inflammation. Thus, IκBζ functions as a molecular switch that reprograms regulatory T cells into Th2-like Treg cells, thereby perturbing peripheral immune tolerance.

**Graphical Abstract:** 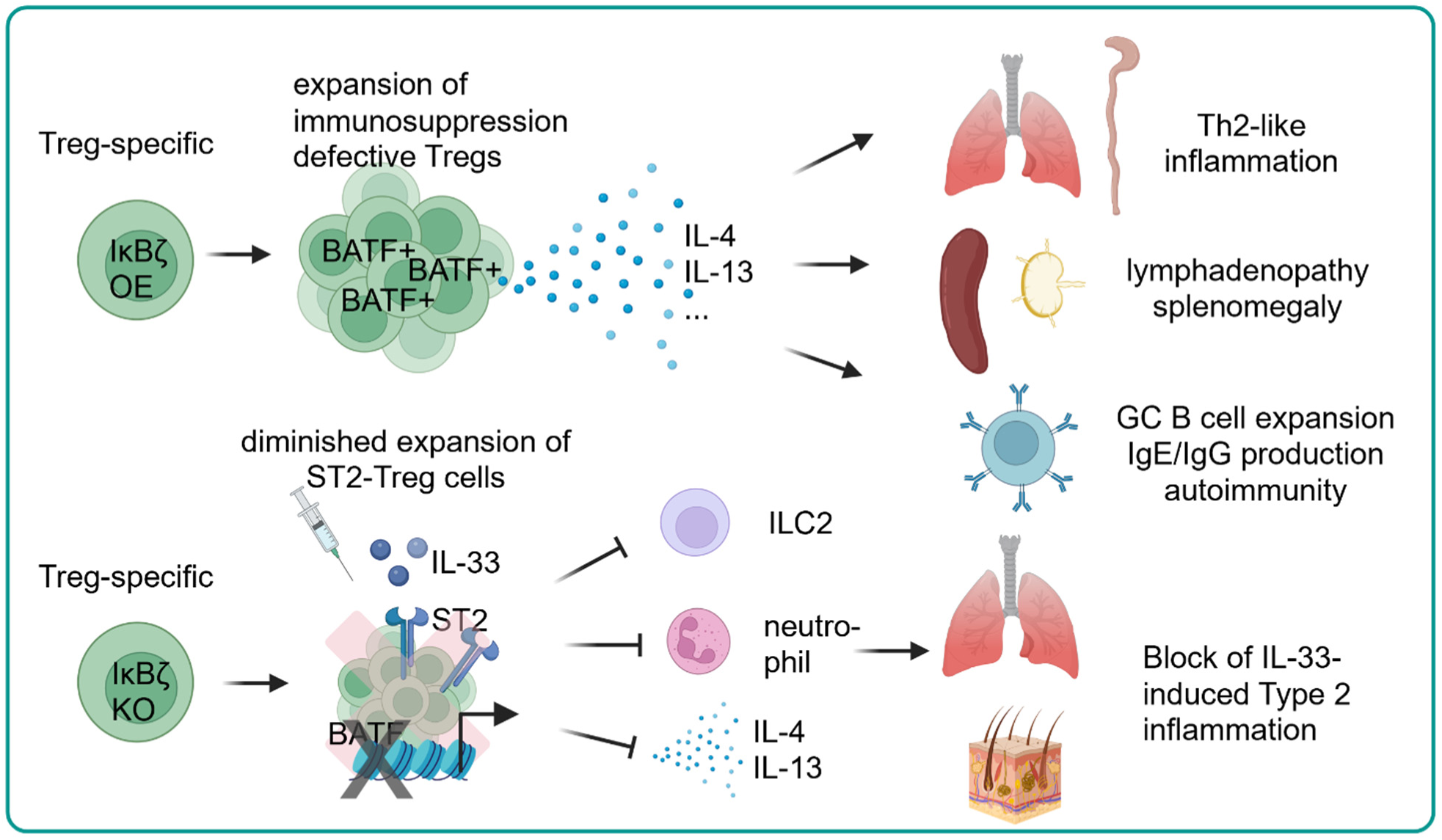

## Introduction

Regulatory T cells constitute a specialized subtype of immune cells that control the balance of inflammatory responses and regulate T and B cell activities, thereby preserving tissue homeostasis and preventing autoimmunity ^1^. Beyond this canonical immunosuppressive role, Treg cells also exert multiple additional organ-specific roles, including tissue repair, metabolic control, and epithelial barrier maintenance ^2, 3^. By the expression of certain cytokines, including IL-10 and amphiregulin, these tissue-resident Treg cells promote tissue regeneration, alleviate tissue damage, and limit fibrosis ^4^.

Tissue-resident Treg cells constitute a heterogeneous subpopulation of FoxP3-positive T cells that are functionally distinct from their lymphoid counterparts and are characterized by increased expression of the IL-33 receptor ST2 ^5^. Upon exposure to IL-33, an alarmin released upon tissue damage, tissue-resident Treg cells can locally expand to limit tissue damage and contribute to tissue homeostasis ^5^. Several transcription factors have been identified that shape tissue-resident Treg function and control their stability and plasticity. For instance, BATF, GATA3, or RBPJ promote or repress the differentiation, homing, and activation of Th2-like Treg cells in certain tissues in mice ^6, 7^. Moreover, key Treg cell-stabilizing factors, such as FoxP3 and TGF-β, suppress effector differentiation and Th2-like features, thereby maintaining the immunosuppressive capacities of Treg cells in lymphoid tissues ^8–10^. How this balance between tissue adaptation and Treg cell lineage stability is molecularly controlled remains incompletely understood, though.

IκBζ, encoded by the *NFKBIZ* gene, is an important cofactor of NF-κB, known to induce or repress a selective subset of NF-κB target genes ^11^. Unlike classical IκB proteins, IκBζ is inducibly expressed in the nucleus of various immune and non-immune cells, including Th17 cells ^12^, NK cells ^13^, macrophages ^14^, and keratinocytes ^15^ upon T cell receptor (TCR) activation, or stimulation with TLR ligands or certain cytokines. Subsequently, IκBζ localizes to chromatin, where it is mandatory for the inducible expression of multiple cytokines and chemokines, including IL-17A, IL-1β, IL-6, IL-36, CXCL1, CXCL2, or CXCL5 ^16^. How IκBζ regulates gene expression is poorly understood. As IκBζ itself lacks a DNA-binding domain, it is thought to act as a bridging factor that recruits epigenetic modifiers to chromatin-bound transcription factors, such as NF-κB or STAT3 ^17, 18^. Notably, aberrant IκBζ expression has been linked to the pathogenesis of multiple inflammatory and autoimmune diseases, including psoriasis ^15, 19^, experimental autoimmune encephalomyelitis (EAE) ^12^, or inflammatory bowel disease ^20^.

Despite its established role in inflammatory gene regulation, the function of IκBζ in Treg cells remains poorly defined. Initial studies using Lck-Cre–mediated deletion of *Nfkbiz* suggested that IκBζ may repress FoxP3 expression ^21^, while Treg cells lacking *Nfkbiz* lost their immunosuppressive function in a T-cell transfer colitis model ^22^. Notably, these defects of IκBζ KO Treg cells were not detectable under steady-state *in vitro* conditions, suggesting that IκBζ may become only functionally relevant in specific inflammatory micromilieus or tissue contexts ^22^. This notion is particularly intriguing because IL-33 signaling, an essential driver of tissue Treg function ^23, 24^, has been reported to induce IκBζ expression in other immune cells, such as mast cells ^25^. Whether IκBζ is a downstream mediator of IL-33 signals in Treg cells, thus controlling tissue Treg differentiation and function, is unknown so far.

In the present study, we investigated the role of IκBζ in Treg cells using Treg-specific gain- and loss-of-function approaches of IκBζ. By generating a Treg-specific IκBζ overexpression mouse model, we show that enforced IκBζ expression drives the expansion of functionally impaired Treg cells with a profound cell-intrinsic Th2-like phenotype, inducing hyperreactive B cell responses and type 2 inflammation in various organs. Mechanistically, IκBζ enhanced the expression and modified the function of BATF, leading to a BATF-dependent expression of Th2-associated cytokines, including IL-4 and IL-13, in the functionally impaired Treg cells. Conversely, Treg-specific deletion of IκBζ in mice impaired BATF function, leading to a diminished expansion of ST2-positive Treg cells upon systemic treatment with IL-33. This led to surprisingly alleviated inflammatory responses in IL-33-treated Treg-specific IκBζ KO mice, due to diminished ILC2 expansion, decreased eosinophilia, and reduced type-2 cytokine expression. Taken together, our findings identify IκBζ as a critical molecular switch that controls Treg cell plasticity, thereby balancing immune tolerance and type 2 inflammation.

## Results

### Generation and validation of Treg-specific IκBζ-overexpressing mice

To investigate the function of IκBζ in Treg cells, we generated a transgenic mouse model enabling the Treg-specific overexpression of IκBζ (OE^ΔTreg^) under the control of a doxycycline (Dox)-inducible system (Figure 1A, Supplementary Figure S1A). In this model, expression of IκBζ (encoded by the *Nfkbiz* gene) is restricted to Foxp3-expressing Treg cells. To validate the functionality of this model, we isolated splenic CD4^+^ T cells and examined IκBζ expression levels in the absence or presence of Dox addition. Compared to control animals lacking Cre recombinase expression, *Nfkbiz* OE^ΔTreg^ mice showed increased *Nfkbiz* mRNA levels in CD25^+^ regulatory T cells. Notably, induction of *Nfkbiz* mRNA occurred independently of the addition of doxycycline (Figure 1B). Furthermore, *in vitro-*differentiated iTreg cells generated from splenic CD4^+^ T-cells, or CD4^+^CD25^+^ sorted splenic Tregs derived from *Nfkbiz* OE^ΔTreg^ mice, showed a strong increase in *Nfkbiz* expression regardless of Dox supplementation (Figure 1C and D). Thus, IκBζ is constitutively expressed in Treg cells in this mouse model, irrespective of Dox administration.

**Figure 1.**
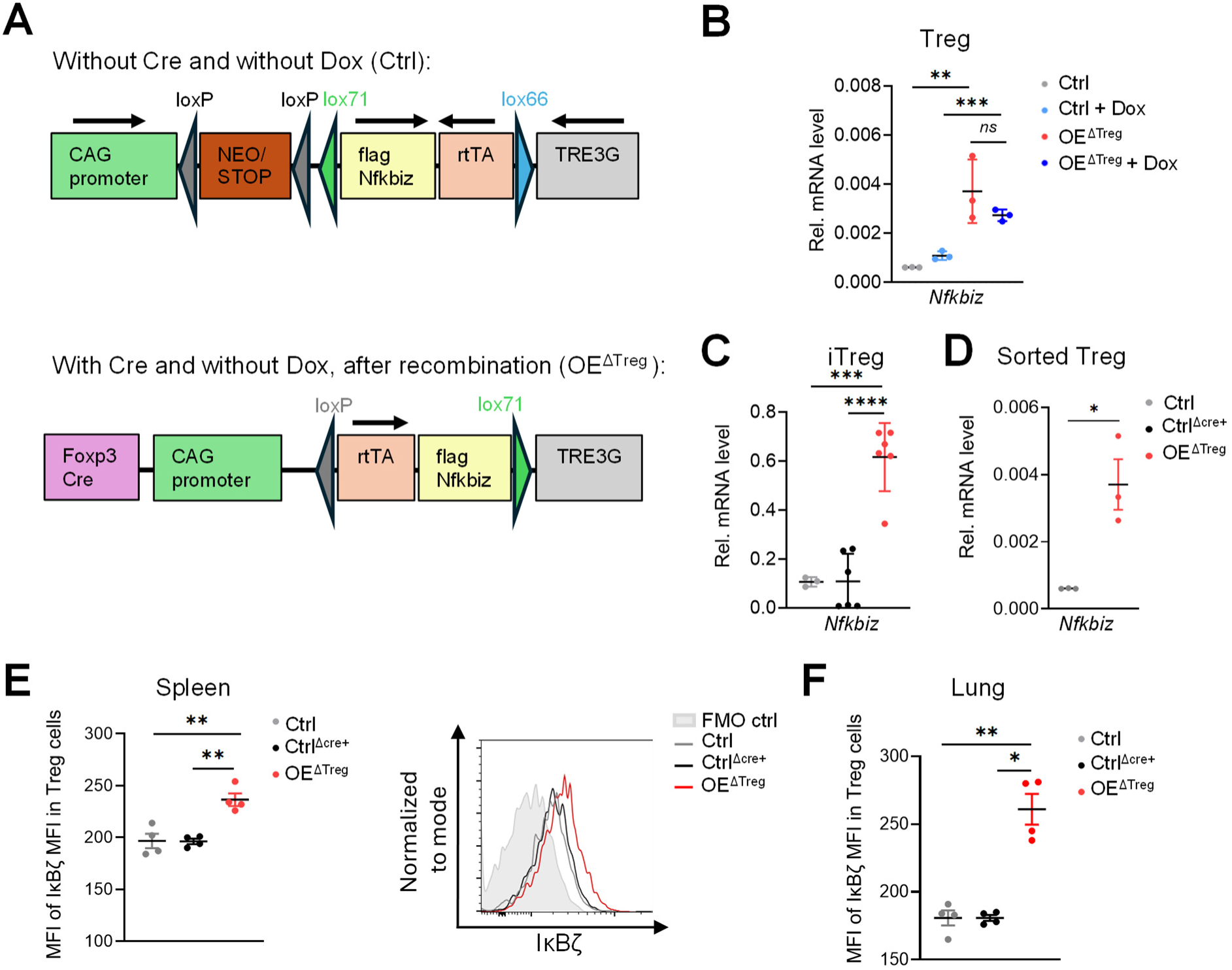
Generation of a mouse model with constitutive overexpression of IκBζ in regulatory T cells. **A.** Schematic illustration of the cloning cassette for the generation of a tissue-specific flag-tagged IκBζ (*Nfkbiz*) overexpression. **B.** *Nfkbiz* mRNA levels in untreated and 72 h doxycycline-treated Treg (CD4^+^ CD25^+^) and conventional T cells (CD4^+^ CD25^-^) isolated from the spleens of Ctrl and OE^ΔTreg^ mice. Shown are the relative mRNA levels of *Nfkbiz* normalized to the housekeeping gene *Actb*. N = 3 independent replicates per group ± SD. **C.** Relative mRNA levels of *Nfkbiz* from *in vitro* differentiated iTreg cells of splenic CD4^+^ T-cells from 12-week-old Ctrl, FoxP3-Cre positive Ctrl (Ctrl^Δcre+^), and Treg-specific IκBζ OE (OE^ΔTreg^) mice, cultured without doxycycline. mRNA levels were normalized to *Actb*. N = 3 - 6 biological replicates per group ± SD. **D.** Results of similar experiments as in panel C, but from CD4^+^ and CD25^+^ FACS-sorted splenic Treg cells of the same mouse strains. Data represent the mean ± SEM. **E + F.** IκBζ protein levels in FoxP3^+^ Treg cells from 10-13-week-old Ctrl, Ctrl^Δcre+,^ and Treg-specific IκBζ-overexpressing mice as detected by flow cytometry. Shown is the mean of 4 biological replicates ± SEM per group. **E.** Data from the spleen of Ctrl, Ctrl^Δcre+,^ and OE^ΔTreg^ mice. *Left:* MFI of IκBζ in Treg cells. *Right:* Example plot of the IκBζ staining normalized to mode, pre-gated on viable, FoxP3^+^ CD4^+^ cells, and a sample that contains all antibodies except IκBζ (FMO) is depicted. **F.** MFI of IκBζ in Treg cells from the lung of Ctrl, Ctrl^Δcre+,^ and OE^ΔTreg^ mice. Significance was calculated using a 2-tailed Student’s t-test (\**p* ≤ 0.05, \*\**p* < 0.01, \*\*\**p* < 0.001, \*\*\*\**p* < 0.0001).

Next, we assessed IκBζ expression in Treg cells at the protein level. Unlike control animals expressing either the Tet-on cassette (Ctrl) or the Foxp3-Cre cassette (Ctrl^Δcre+^) alone, FoxP3-positive Treg cells from *Nfkbiz* OE^ΔTreg^ mice showed a significantly increased expression of IκBζ in Treg cells from the spleen (Figure 1E) and lung (Figure 1F). Thus, as constitutive IκBζ expression was already present without the additional treatment with Dox, and additional Dox treatment might lead to potential off-target effects, subsequent experiments were performed without Dox supplementation to specifically assess the consequences of Treg-specific overexpression of IκBζ.

### Overexpression of IκBζ in Treg cells leads to an expansion of potentially dysfunctional Treg cells, resulting in lymphadenopathy and splenomegaly

To characterize the consequences of IκBζ overexpression on Treg cell numbers and function, we examined secondary lymphoid organs, such as spleen and lymph nodes, as well as non-lymphoid organs, such as the lung and colon, in young (11 - 16-week-old) and aged (≥ 30 weeks old) animals. Already at the macroscopic level, slightly enlarged lymph nodes and spleens could be detected in young animals (Figure 2A), which progressed to severe lymphadenopathy and splenomegaly in aged animals (Figure 2B). Notably, this enlargement was associated with an increased proportion of Treg cells in secondary lymphoid organs (Figure 2C) and other organs, such as the lung and colon (Figure 2D), but surprisingly not in the skin (Figure 2E). A comparable expansion of Treg cells could also be detected in aged animals (Supplementary Figure S2A and B). Of note, enforced expression of IκBζ did not perturb FoxP3 expression in Treg cells. Instead, we detected a modest decrease in relative FoxP3 expression in both the Ctrl^Δcre+^ and OE^ΔTreg^ strains when compared with the Ctrl, due to the use of the Sakaguchi Foxp3-cre strain (Supplementary Figure S2C).

**Figure 2.**
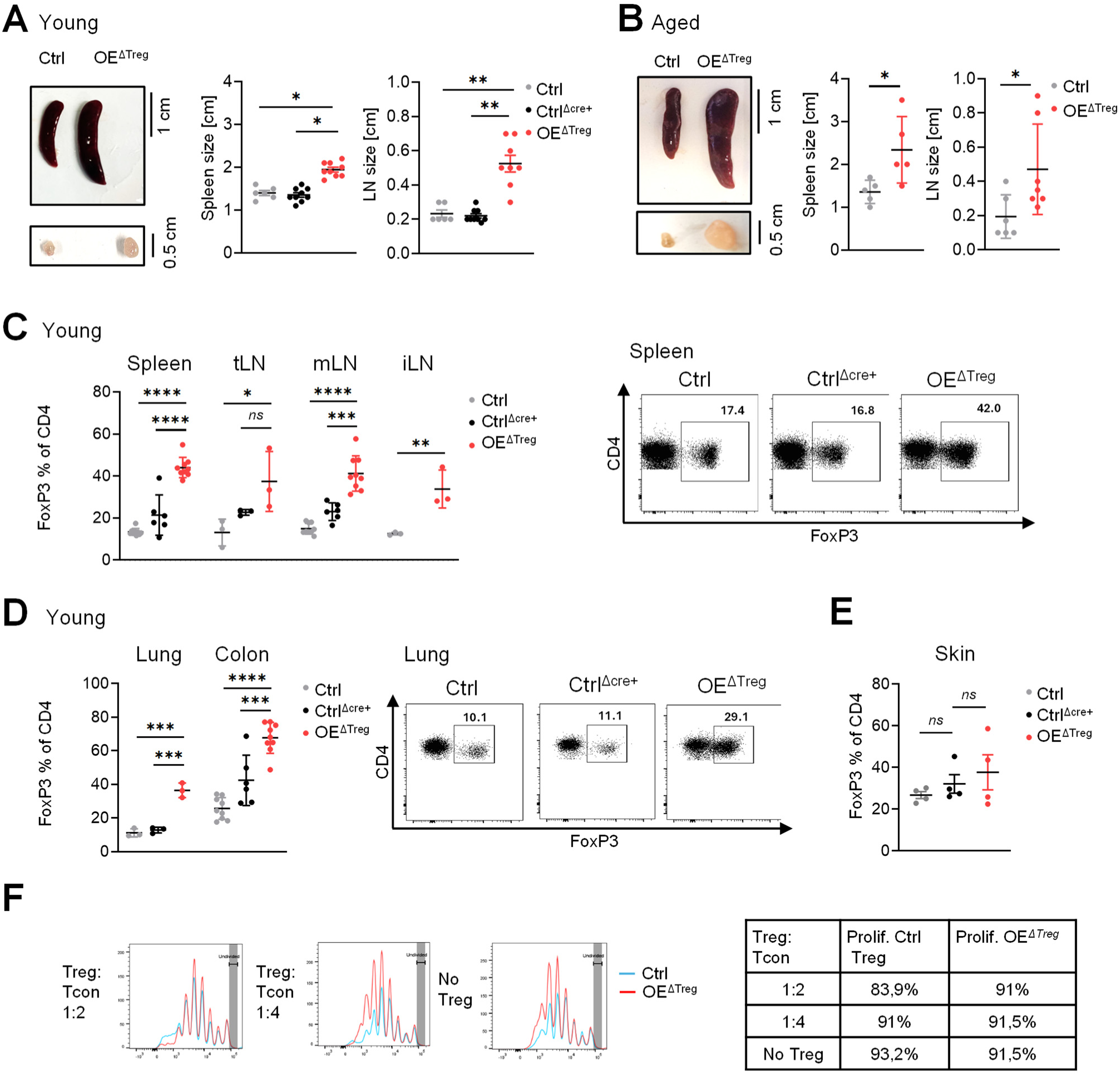
Treg-specific overexpression of IκBζ causes a massive expansion of immunosuppression-defective Treg cells in various organs. A +. **B.** Representative pictures and size of spleen and lymph nodes from Ctrl, Ctr^Δcre+^, and *Nfkbiz* OE^ΔTreg^ mice. **A.** Young mice, around 11-16-weeks old. N = 6 – 9 mice per group ± SEM. **B.** Aged mice, ≥ 30-weeks old. N = 5 – 7 mice per group. **C - E.** Flow cytometric analysis of FoxP3^+^ Treg cells within living CD4^+^ cells from 11-16-week-old mice. **C.** Percentage of FoxP3^+^ Treg cells within living CD4^+^ cells of spleen and lymph nodes. tLN = tracheal lymph node, mLN = mesenteric lymph node, iLN = inguinal lymph node. *Right site:* Representative dot plots of FoxP3 staining, illustrated over CD4, pre-gated on living CD4^+^ cells. N = 3 – 9 mice per group ± SEM. **D.** Percentage of FoxP3^+^ Treg cells within living CD4^+^ cells in the lung and colon, and representative dot plots showing FoxP3 and CD4 staining of the cells, pre-gated on living CD4^+^ cells. N = 3 – 9 mice per group ± SEM. **E.** Percentage of FoxP3^+^ Treg cells within living CD4^+^ cells of the skin. N = 4 mice per group ± SEM. F. *In vitro* immunosuppression assay of Ctrl and *Nfkbiz*-overexpressing Treg cells. Significance was calculated using a 2-tailed Student’s t-test (\**p* ≤ 0.05, \*\**p* < 0.01, \*\*\**p* < 0.001, \*\*\*\**p* < 0.0001, *ns* = not significant).

As a major function of Treg cells relies on their ability to suppress effector T-cell proliferation, we next tested whether IκBζ-overexpressing Treg cells retained their immunosuppressive capacity. A classical *in vitro* immunosuppression assay using wild-type CD4^+^ CD25^-^ responder T cells and Treg cells from *Nfkbiz* OE^ΔTreg^ or Ctrl^Δcre+^ mice showed that IκBζ-overexpressing Treg cells had only reduced suppressive capacity compared with Ctrl^Δcre+^ Treg cells (Figure 2F). Taken together, IκBζ overexpression in Treg cells induces their expansion in several organs. At the same time, these Treg cells might become dysfunctional, at least with respect to their ability to repress effector T-cell proliferation.

### Treg-specific overexpression of IκBζ induces a Th2-associated disease pattern

As IκBζ-overexpressing Treg cells partially lost their immunosuppressive function, at least *in vitro*, we next sought to investigate the pathological consequences *in vivo*, beyond the severe lymphadenopathy and splenomegaly observed in the aged animals. We focused especially on “barrier” organs, which are constantly exposed to microorganisms and pathogen-associated molecular patterns and therefore critically rely on a balanced tissue immune homeostasis.

Unlike in secondary lymphatic organs, no macroscopic abnormalities were observed in the colon and skin of young *Nfkbiz* OE^ΔTreg^ animals (Supplementary Figure 3A). However, in aged mice, we detected an increased prevalence of spontaneously arising dermatitis (Supplementary Figure S3B and C), as well as a shortened colon length, indicative of colon inflammation (Supplementary Figure S3D). Furthermore, a detailed histological examination of the lung revealed a modest influx of inflammatory infiltrates in young *Nfkbiz* OE^ΔTreg^ mice, which dramatically increased in older animals (Figure 3A). This phenotype was accompanied by a significant expansion of mucus-producing goblet cells in the airways of young and aged Treg-specific IκBζ-overexpressing mice (Figure 3B), as well as increased collagen deposition, indicative of fibrosis induction (Supplementary Figure S3E).

**Figure 3.**
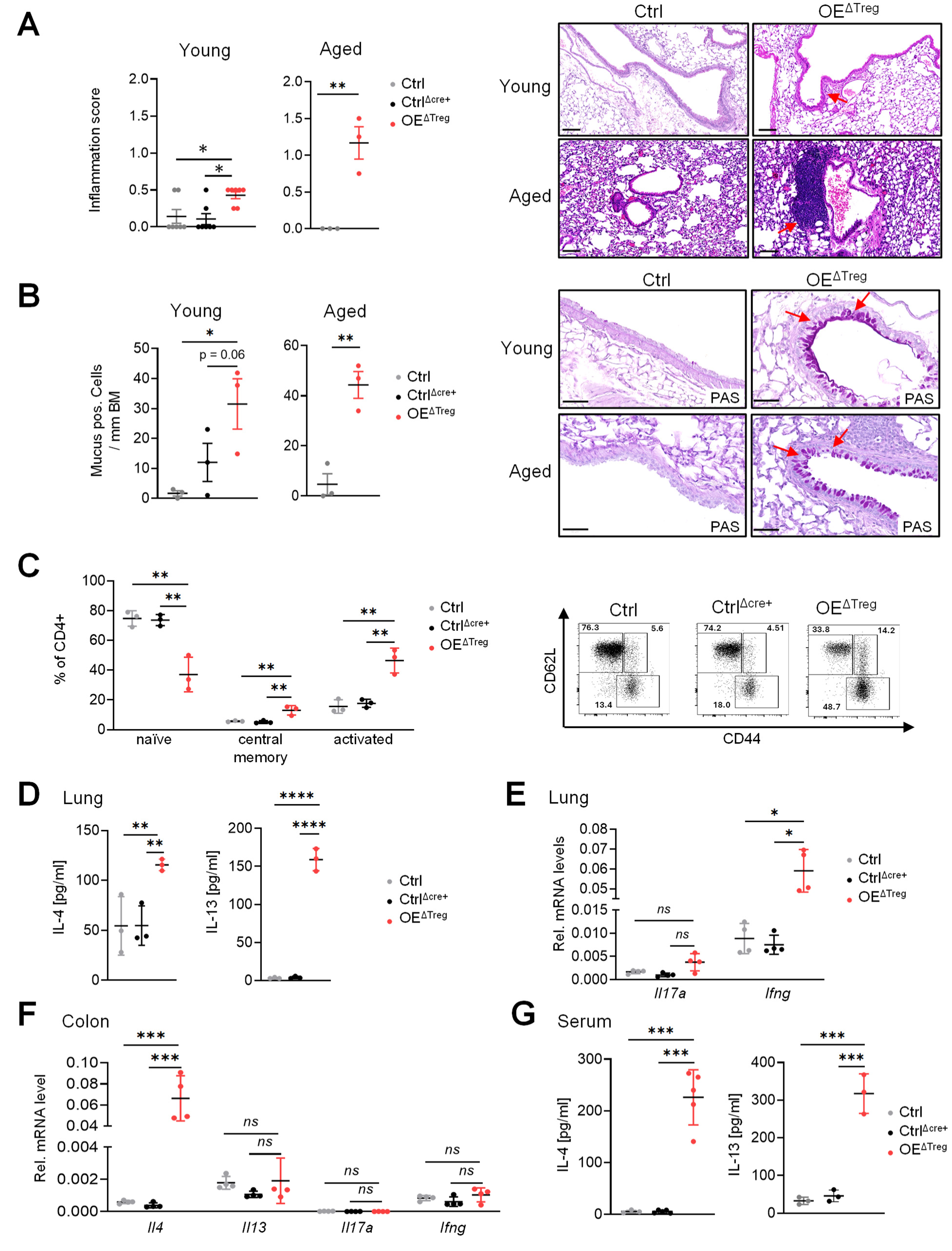
Overexpression of IκBζ in Treg cells causes systemic type 2 inflammation. **A.** Inflammation score reflecting the relative amount of immune cell infiltrates in the lung of young and aged Treg-specific IκBζ overexpressing mice. Representative images are shown on the right. Scale: 100 µm. N = 3 - 7 mice per group. **B.** Relative amount of mucus-producing goblet cells, per mm basal membrane (BM), counted from periodic acid Schiff (PAS)-stained lungs of young and old Ctrl, Ctr^Δcre+^, and *Nfkbiz* OE^ΔTreg^ mice. Representative images are shown on the right. Scale: 50 µm. N = 3 mice per group. **C.** Flow cytometry analysis of effector T cells in the lung of 11-13-week-old mice. Cells were pre-gated on living CD3^+^, CD4^+^ cells. Naïve T-cells = CD62L^+^ CD44^-^, central memory T-cells = CD62L^+^ CD44^+^, activated T-cells = CD62L^-^ CD44^+^. N = 3 mice per group ± SEM. Right site: representative dot plots of all three experimental groups. **D.** IL-4 and IL-13 protein levels in lung tissue of young mice, measured by ELISA. N = 3 mice per group ± SEM. **E.** Gene expression of *Il17a* and *Ifng* in lung tissue of young mice, normalized to *Actb*. N = 4 mice per group ± SEM. **F.** Gene expression analysis of colon tissue from aged mice (> 30 weeks old), normalized to *Actb*. N = 4 mice per group ± SEM. **G.** Protein levels of IL-4 and IL-13, measured by ELISA, in the plasma of aged mice. N = 4 mice per group ± SEM. Significance was calculated using a 2-tailed Student’s t-test (\**p* ≤ 0.05, \*\**p* < 0.01, \*\*\**p* < 0.001, \*\*\*\**p* < 0.0001, *ns* = not significant).

As the lung, compared to the colon and skin, showed the earliest and most significant abnormalities in the Treg-specific IκBζ-overexpressing mice, we further characterized the cellular and molecular characteristics underlying these pathological changes. Notably, goblet cell metaplasia and increased presence of mucus are tightly linked to the local expression of Th2-derived cytokines, such as IL-4 and IL-13 ^26^. In agreement with this hypothesis, young *Nfkbiz* OE^ΔTreg^ mice already exhibited an increased proportion of CD44^+^/CD62L^-^ activated T cells in the lung, whereas the relative number of naïve T cells decreased (Figure 3C). The relative increase in activated T cells and simultaneous decrease in naïve T cells were even more pronounced in aged *Nfkbiz* OE^ΔTreg^ mice (Supplementary Figure S3F). Furthermore, this activation of effector T cells was accompanied by a strong increase in the expression of IL-4 and IL-13 in the lung tissue of young mice (Figure 3D), whereas *Il17a* was hardly detectable, and *Ifng* expression was only mildly increased in the lungs of *Nfkbiz* OE^ΔTreg^ mice (Figure 3E).

Based on these results, we hypothesized that the observed skin and colon inflammation in the aged *Nfkbiz* OE^ΔTreg^ mice also resembles a dominant Th2 signature. As expected, gene expression analysis of Th1-, Th2-, and Th17-associated cytokines revealed a strong Th2 polarization in inflamed skin areas and colon tissues of aged *Nfkbiz* OE^ΔTreg^ mice, marked by an elevated expression of *Il4* (Figure 3F, Supplementary Figure S3G). Furthermore, we detected a strong increase in IL-4 and IL-13 protein levels in the serum of aged Treg-specific IκBζ-overexpressing mice, indicating a systemic shift towards Th2 inflammation (Figure 3G). In conclusion, Treg-specific overexpression of IκBζ impairs Treg function, leading to uncontrolled Th2-driven inflammation, which especially affects barrier organs such as the lung.

### IκBζ overexpression in Treg cells leads to hyperactive B cells marked by IgE class switch and a massive production of autoreactive antibodies

Th2 cytokines induce a class switch in B cells, leading to IgE antibody production ^27^. As we detected an increased systemic expression of the Th2 cytokines IL-4 and IL-13 in *Nfkbiz* OE^ΔTreg^ mice, we wondered whether B cells were also affected by the Treg-specific overexpression of IκBζ. We therefore investigated potential changes in B cell prevalence, activity, and effector responses in the spleen, lymph nodes (LN), and the lung. Interestingly, the total number of B cells was only mildly increased in young *Nfkbiz* OE^ΔTreg^ mice (Figure 4A). However, we detected an increased activation of B cells in the tracheal LN and the lung, shown by elevated MHC class II expression in young mice (Figure 4B, left), with a further increase in aged *Nfkbiz* OE^ΔTreg^ mice (Figure 4B, right). At the same time, we found an increase of CD95^+^CD38^-^ germinal center (GC) B cells in the spleen and lymph nodes of young and aged mice with Treg-specific IκBζ overexpression, indicating elevated antibody production (Figure 4C).

**Figure 4.**
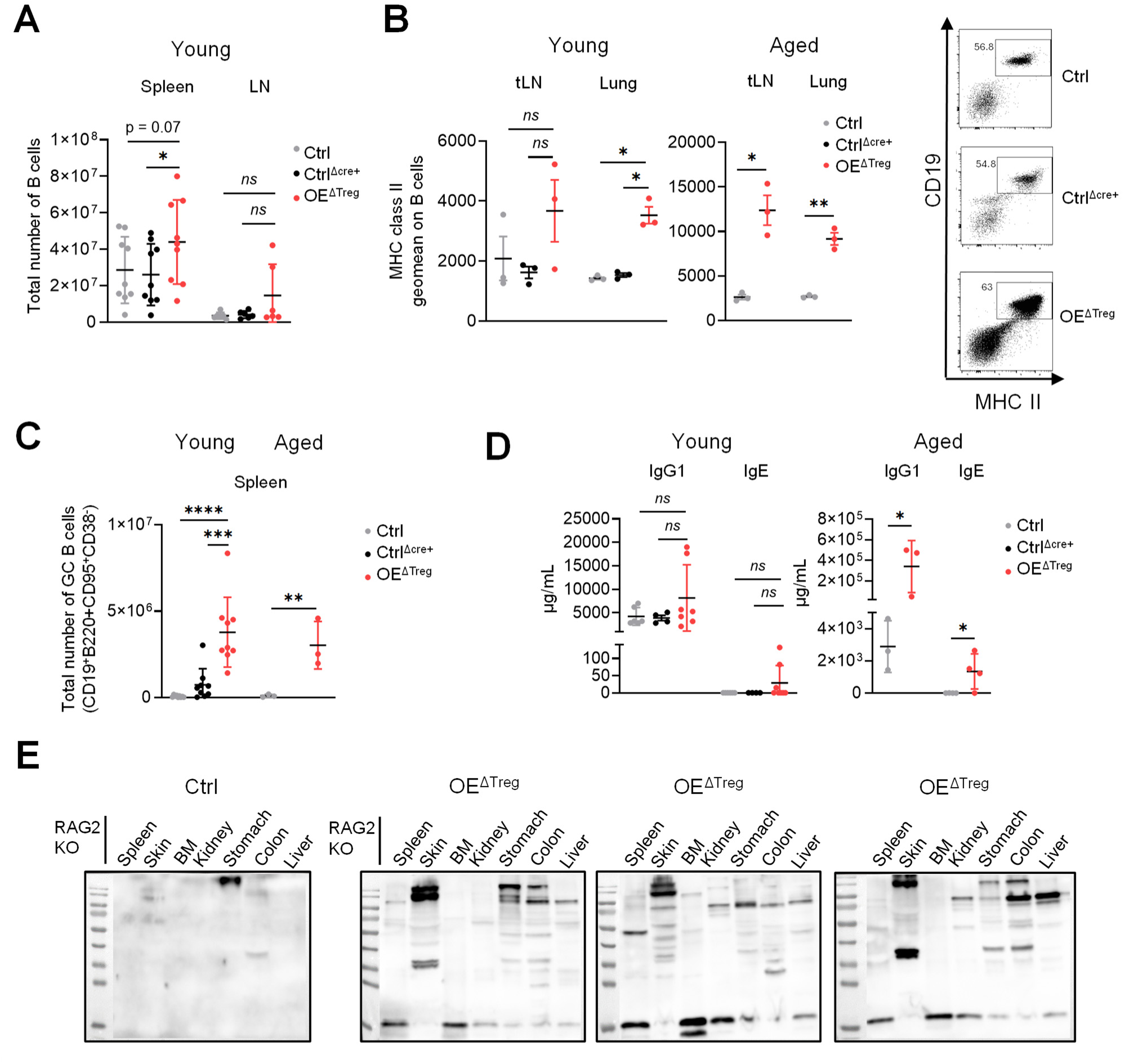
Overexpression of IκBζ in Treg cells induces B-cell activation, GC B-cell expansion, and the increased production of IgE and IgG antibodies. **A.** Total number of living CD19^+^ B cells in the spleen and lymph node of young mice (11-16 weeks old). **B.** Geomean of MHC class II expression on CD19^+^ B-cells in tracheal lymph nodes (tLN) and the lung of young and aged mice. N = 3 mice per group ± SEM. *Right*: Representative histograms of MHCII expression on B cells. **C.** Total number of GC B cells in the spleen of young and aged (≥ 30-weeks-old) mice, identified as viable CD19^+^, B220^+^, CD95^+^, CD38^-^ cells. N = 3 - 8 mice per group ± SEM. **D.** IgG1 and IgE plasma levels in young and aged mice, determined by ELISA. N = 3 – 6 mice per group ± SEM. **E.** Validation of the presence of autoreactive antibodies using plasma from aged control and Treg-specific IκBζ-overexpressing mice. Protein lysates were generated from various tissues of RAG2 knockout (KO) mice, which were equally loaded and analysed by Western blotting. Serum samples of one control and 3 *Nfkbiz* OE^ΔTreg^ mice were used as primary antibodies. Immunoreactive protein bands indicate the presence of autoreactive antibodies in the serum of Treg-specific IκBζ-overexpressing mice. Significance was calculated using a 2-tailed Student’s t-test (\**p* ≤ 0.05, \*\**p* < 0.01, \*\*\**p* < 0.001, \*\*\*\**p* < 0.0001, *ns* = not significant).

To verify whether the enhanced B cell activation phenotype was associated with increased antibody secretion, we analyzed antibody titers in the sera of young and aged mice. Notably, IgG1 antibodies were slightly elevated in young animals, whereas IgE was not detectable. However, dramatically elevated titers of IgG1 and IgE antibodies were present in the sera of aged *Nfkbiz* OE^ΔTreg^ mice compared to controls (Figure 4D). The IgG1 antibodies resembled autoantibodies, as detected by immunoblot analysis of tissue samples from T- and B-cell-deficient RAG2 knockout mice (Figure 4E). In detail, we incubated the membranes containing blotted protein lysates from various organs of RAG2 knockout mice with the serum of Ctrl and *Nfkbiz* OE^ΔTreg^ mice. In contrast to serum from control mice, which rendered no significant protein bands, the serum of three different aged *Nfkbiz* OE^ΔTreg^ mice was autoreactive and detected a wide range of endogenous proteins from various organs (Figure 4E). In summary, this data implies that the malfunction of Treg cells harboring IκBζ overexpression results in a global hyperactivity of B cells. With increasing age, *Nfkbiz* OE^ΔTreg^ mice exhibit the production of autoreactive IgG and IgE antibodies, reflecting the onset of autoimmunity in the context of systemic Th2-driven inflammation.

### Intrinsic expression of Th2-like cytokines in Treg-specific IκBζ-overexpressing mice

So far, we have demonstrated a partial loss of the immunosuppressive function of IκBζ-overexpressing Treg cells, accompanied by a systemic Th2 polarization leading to B cell class switching and lung inflammation. However, it remained unclear whether these changes are triggered intrinsically in Treg cells or represent secondary effects in effector T cells due to impaired Treg cell function. To dissect the cell-intrinsic effects of Treg-specific IκBζ overexpression, we performed single-cell sequencing of CD45^+^ cells from spleens from young and aged *Nfkbiz* OE^ΔTreg^ mice and corresponding age- and gender-matched control animals (harboring only the *Nfkbiz*-encoding Tet-on cassette) (Figure 5A, Supplementary Figure S4A and B). Similar to the previous flow cytometry analysis, we detected an expansion of two distinguishable Treg subclusters, which we, based on a previous publication ^5^, termed naïve Treg and tissue-resident Treg cells (Figure 5A and B, Supplementary Figure S4B-D). Furthermore, we detected decreased numbers of naïve and activated CD8^+^ T cells, whereas GC B cells, and in the aged mice, also monocytes and macrophages, increased. Interestingly, when we further investigated gene expression changes in these two Treg cell clusters, we found a strong upregulation of Th2-associated key molecules, such as *Il13*, *Il4*, and *Batf,* within the Treg cell clusters (Figure 5C and D). These gene expression changes could further be validated in FACS-sorted, CD4^+^CD25^+^, IκBζ-overexpressing Treg cells from the spleen (Figure 5E). Collectively, this data implies that IκBζ overexpression in Treg cells induces a Th2-like gene expression program intrinsically in Treg cells. This phenotype likely reflects a selective expansion or functional reprogramming of tissue-resident Treg cells toward Th2-like effector functions, thereby contributing to the systemic immune dysregulation in *Nfkbiz* OE^ΔTreg^ mice.

**Figure 5.**
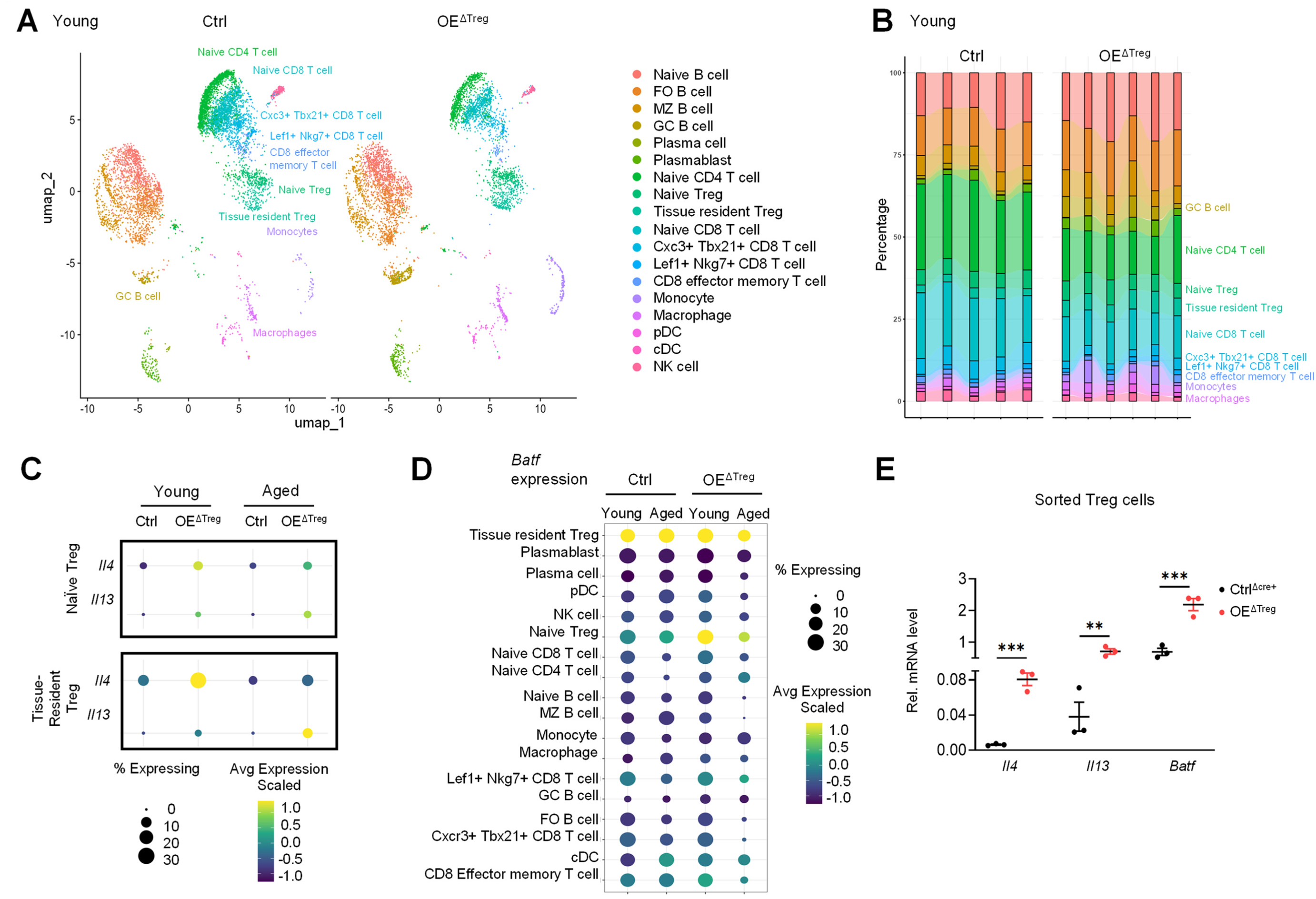
Single-cell sequencing of the spleen identifies an expansion of BATF-positive, type 2 cytokine-expressing Treg cells in *Nfkbiz* OE^ΔTreg^ mice. **A.** UMAP plot of the identified cell types in the spleen of 11-week-old control and Treg-specific IκBζ-overexpressing mice. **B.** Relative numbers and distribution of the identified cell type clusters in all sequenced mice. Spleens of 3 control and 3 *Nfkbiz* OE^ΔTreg^ mice were sequenced in duplicate. One control sample did not pass the quality assessment and was not further evaluated. Shown is the percentage of the respective cell type relative to the total number of sequenced cells. Markers for the cluster annotation can be retrieved from Supplementary Figure S4A and B. **C.** Dot plots showing the average expression levels of *Il4* and *Il13,* and the relative number of positive cells within the Treg clusters identified by single-cell sequencing of spleens from young and middle-aged mice. Notably, the specific markers defining the Treg clusters were derived from a previous publication ^5^ (naïve Treg: CCL5^high^; tissue-resident Treg cells: Tigit^high^ Ikzf2^high^, Nrp1^high^). D. *Batf* expression levels and the relative number of positive cells across all annotated cell types were identified by single-cell sequencing of young and middle-aged mice. **E.** Gene expression analysis in CD4^+^ CD25^+^-sorted splenic Tregs from control and *Nfkbiz* OE*^ΔTreg^* mice. N = 3 mice per group ± SEM. Significance was calculated using a 2-tailed Student’s t-test (\*\**p* < 0.01, \*\*\**p* < 0.001).

### IκBζ overexpression in Treg cells promotes a BATF-dependent differentiation into Th2-like tissue-resident Treg cells

To test our hypothesis that IκBζ regulates a Th2-like tissue Treg program, we performed an unbiased transcriptome profiling of *in vitro-*differentiated Treg (iTreg) cells from splenic CD4^+^ T cells of control and *Nfkbiz* OE^ΔTreg^ mice. Upon IκBζ overexpression, 57 genes were significantly downregulated, and 320 genes were significantly upregulated in iTreg cells. The top upregulated genes included markers associated with Treg activation, such as *Icos* and *Ctla4*, but also markers representing a previously published signature of tissue-resident Treg cells, including *Ebi3*, *Batf, Il4, Il13, Il1rl1,* and *Klrg1* ^28^ (Figure 6A, Supplementary Figure S5A). Of note, expression of key molecules, such as *Foxp3* and *Tgfb1*, which were previously shown to restrain Th2-like features in Treg cells, remained unperturbed or even increased in IκBζ-overexpressing Treg cells (Supplementary Figure S5B).

**Figure 6.**
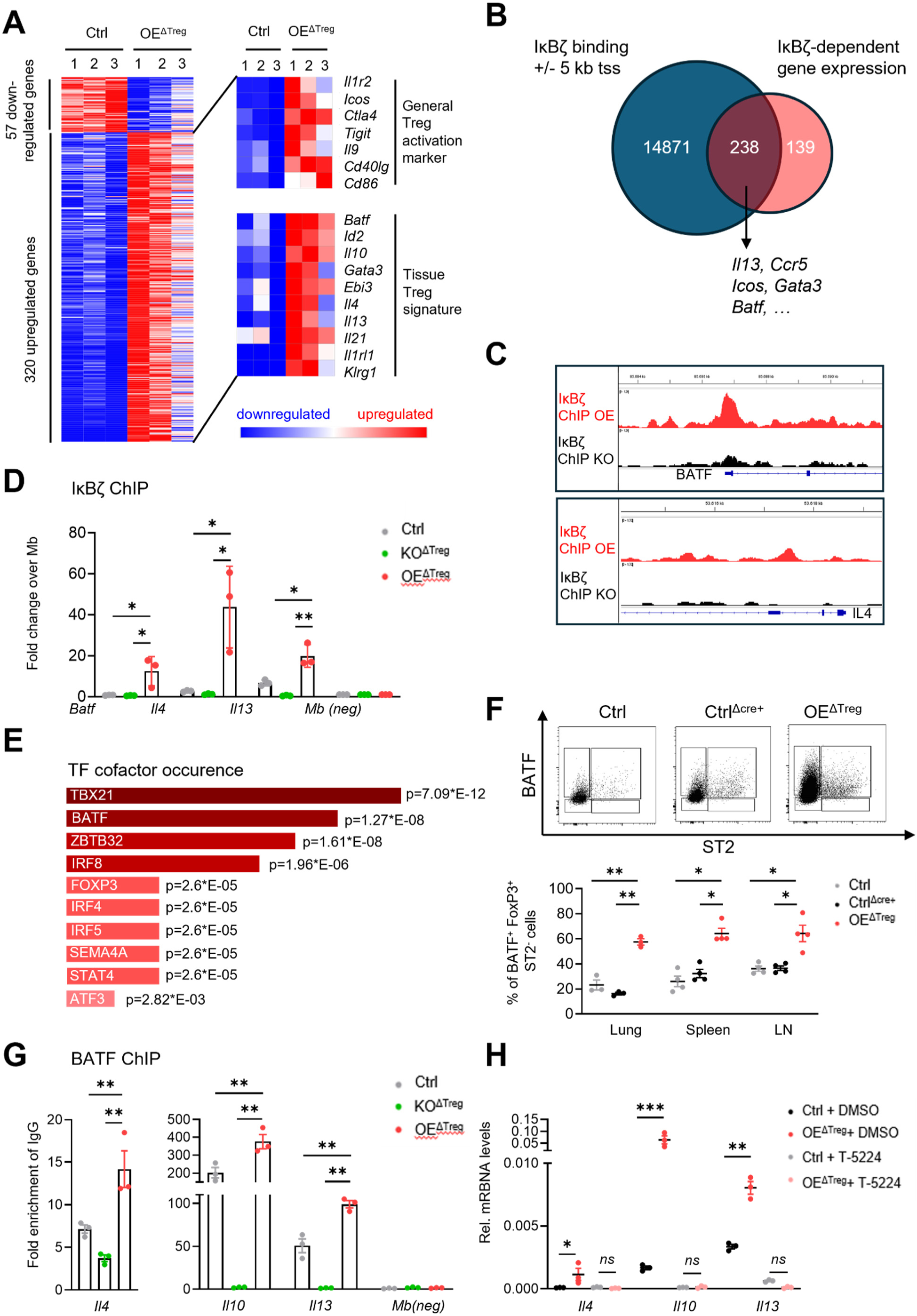
IκBζ induces the expression of Th2-associated cytokines in Treg cells via modulation of BATF expression and function. **A.** Transcriptome analysis of *in-vitro* differentiated control and IκBζ-overexpressing iTreg cells. iTreg cells were differentiated from splenic CD4^+^ positive cells for 4 days. Depicted is a heatmap comprising all differentially regulated genes, calculated using the proportion-based Baggerly test (minimal fold change > 2; Difference > 4; p-value ≤ 0.05). *Right:* heatmap of selected deregulated genes comprising general markers for Treg activation and tissue-resident Treg cells. Red = upregulated, blue = downregulated genes. **B.** Overlay of IκBζ-binding sites and differentially expressed genes (as depicted in A) in iTreg cells. IκBζ-binding sites identified by peak calling from IκBζ ChIP sequencing of iTreg cells with either overexpression (IκBζ ChIP OE) or deletion (IκBζ ChIP KO) of IκBζ. Only binding sites 5 kb up- or downstream of the transcription start site (tss) were considered. **C.** Graphical illustration of the IκBζ ChIP sequencing results at the *Il4* and *Batf* loci. For each gene locus, the maximal height of the tracks was normalized. **D.** Targeted ChIP analysis of IκBζ in control, IκBζ-overexpressing, and IκBζ KO iTreg cells. Depicted is the fold enrichment calculated over the negative control locus *Mb*. N = 3 technical replicates per experimental group. **E.** Transcription factor co-occurrence of IκBζ-binding sites 5 kb up- and downstream of the tss (similar as described in B) was retrieved using Enrichr ^59^. The length of the bars indicates the relative number of IκBζ-bound genes that also contain binding sites for the depicted transcription factors. **F.** BATF protein levels determined by intracellular flow cytometric staining in viable FoxP3^+^ and ST2^-^ cells from spleen, inguinal lymph nodes, and lung of young Ctrl, Ctrl^Δcre^, and *Nfkbiz* OE^ΔTreg^ cells. BATF levels in FoxP3^+^ and ST2^+^ cells remained unchanged (Supplementary Figure S5D). N = 4 mice per group ± SEM. **G.** ChIP analysis of BATF in PMA- and ionomycin (P/I)-stimulated iTreg cells from control, IκBζ OE^ΔTreg^, and IκBζ KO^ΔTreg^ mice. Cells were stimulated for 4 h before ChIP. Shown is the fold enrichment over an IgG control ChIP in IκBζ OE iTregs. The *Mb* gene locus served as a negative control locus. N = 3 biological replicates per group ± SD. **H.** Relative gene expression in Ctrl and IκBζ-overexpressing iTreg cells, stimulated for 24 h with DMSO as vehicle control, or with 10 µM of the JUN/BATF inhibitor T5224. Relative mRNA levels were normalized to *Actb*. N = 3 biological replicates ± SD. Significance was calculated using a 2-tailed Student’s t-test (\**p* ≤ 0.05, \*\**p* < 0.01, \*\*\**p* < 0.001, \*\*\*\**p* < 0.0001, *ns* = not significant).

To dissect which of these deregulated target genes are directly controlled by IκBζ, we performed chromatin immunoprecipitation (ChIP) sequencing of IκBζ using IκBζ-overexpressing iTreg cells. Specificity of the ChIP was controlled by precipitating IκBζ-bound chromatin in IκBζ knockout iTreg cells. Interestingly, the ChIP sequencing analysis revealed that IκBζ localized to 15109 genes, predominantly to the regions 5 kb up- or downstream of the transcription start site (Figure 6B, Supplementary Figure S5C). Furthermore, the majority of potential IκBζ target genes identified by bulk RNA sequencing were directly bound by IκBζ (Figure 6B and C). This was especially the case for type-2 inflammation-related genes, such as *Batf*, *Il4*, and *Il13*, which we could also validate by targeted ChIP of IκBζ in iTreg cells (Figure 6D, Supplementary Figure S5D).

As IκBζ cannot directly bind to chromatin but rather influences the binding of other transcription factors, such as NF-κB, we next investigated which transcription factor-binding sites were enriched in genes directly bound and regulated by IκBζ. Interestingly, analysis of transcription factor co-occupancy identified an enrichment for BATF-binding sites, amongst others, within the pool of direct IκBζ target genes in Treg cells ^29^ (Figure 6E). Notably, BATF was previously shown to drive the differentiation of non-lymphoid tissue Treg precursor cells ^5^. In T follicular helper cells, BATF is known to bind to the *Il4* promoter, triggering its expression ^30^. Given our RNA and ChIP sequencing results, we therefore hypothesized that IκBζ regulates tissue Treg-related gene expression by directly promoting BATF expression and function in Treg cells, especially as *Batf* has been identified as a potential IκBζ target gene, before ^31^. Indeed, Treg-specific overexpression of IκBζ increased BATF expression (Figure 6F). This was, however, only the case in conventional ST2-negative Treg cells, but not in classical tissue-resident ST2-positive counterparts (Supplementary Figure S5E and S5F). Furthermore, Treg-specific overexpression of IκBζ led to increased binding of BATF to the promoter regions of *Il4*, *Il10*, and *Il13* in activated iTreg cells, whereas the deletion of IκBζ completely ablated the binding of BATF to these promoter regions (Figure 6G). Finally, inhibition of BATF using the BATF/JUN inhibitor T-5224 completely reversed the increased gene expression of *Il4*, *Il10*, and *Il13* in IκBζ-overexpressing Treg cells (Figure 6H), thereby establishing BATF as a key transcription factor modulated by IκBζ in Treg cells. Taken together, these findings suggest that constitutive IκBζ expression modulates the expression and function of BATF in Treg cells. This, in turn, leads to an intrinsic induction of Th2-like cytokines, such as *Il4* and *Il13,* in Treg cells, promoting Th2-driven inflammation in several organs, including the lung and colon.

### Treg-specific KO of IκBζ restrains IL-33-driven expansion and Th2-like conversion of Treg cells

Based on our finding that enforced expression of IκBζ in Treg cells modulates BATF function and subsequent expansion of tissue-resident Treg cells, we wondered whether Treg-specific deletion of IκBζ limits the differentiation and function of ST2^+^ Treg cells, possibly by constraining BATF function. To address this question, we generated Foxp3-cre *Nfkbiz* knockout (KO^ΔTreg^) mice by crossing *Nfkbiz* flox Ctrl mice (subsequently used as control) with FoxP3-cre-expressing mice ^32^. Phenotypically, Treg-specific *Nfkbiz* knockout (KO) mice bred normally and did not show signs of a perturbed homeostasis (Supplementary Figure S6A). Detailed flow cytometry analysis revealed mostly unperturbed levels of ST2^+^ Treg cells in the KO^ΔTreg^ mice, with the notable exception of the skin, where ST2^+^ Treg cells were significantly diminished in *Nfkbiz* KO^ΔTreg^ mice compared to control *Nfkbiz* flox mice (Supplementary Figure S6B).

As deletion of *Nfkbiz* in Treg cells did not largely affect steady-state levels of ST2^+^ Treg cells, we next tested if this might be different in a setting of an acute expansion of ST2^+^ Treg cells. Previously, it was shown that the repeated systemic administration of IL-33, the ligand of the ST2 receptor, induces an acute expansion of ST2^+^ tissue-resident Treg cells ^33, 34^. Furthermore, *in vitro* treatment of mast cells with IL-33 was reported to induce IκBζ expression ^25^. We therefore hypothesized that Treg-derived IκBζ might preferentially promote the acute expansion of ST2^+^ Treg cells under inflammatory, but not steady-state conditions. To test this hypothesis, we repeatedly treated control and Foxp3-cre *Nfkbiz* KO mice with IL–33 and examined IκBζ expression in Treg cells, as well as ST2^+^ Treg cells expansion in multiple organs (Figure 7A). As expected, acute systemic IL-33 treatment increased the number of IκBζ-positive ST2^+^ Treg cells, which was not detectable for ST2^-^ Treg cells (Figure 7B). Furthermore, in agreement with previous reports, IL-33 treatment also induced a massive expansion in ST2^+^ Treg cells in the lung, skin, spleen, and lymph nodes. However, this expansion was significantly diminished upon IL-33 treatment of *Nfkbiz* KO^ΔTreg^ mice (Figure 7C).

**Figure 7.**
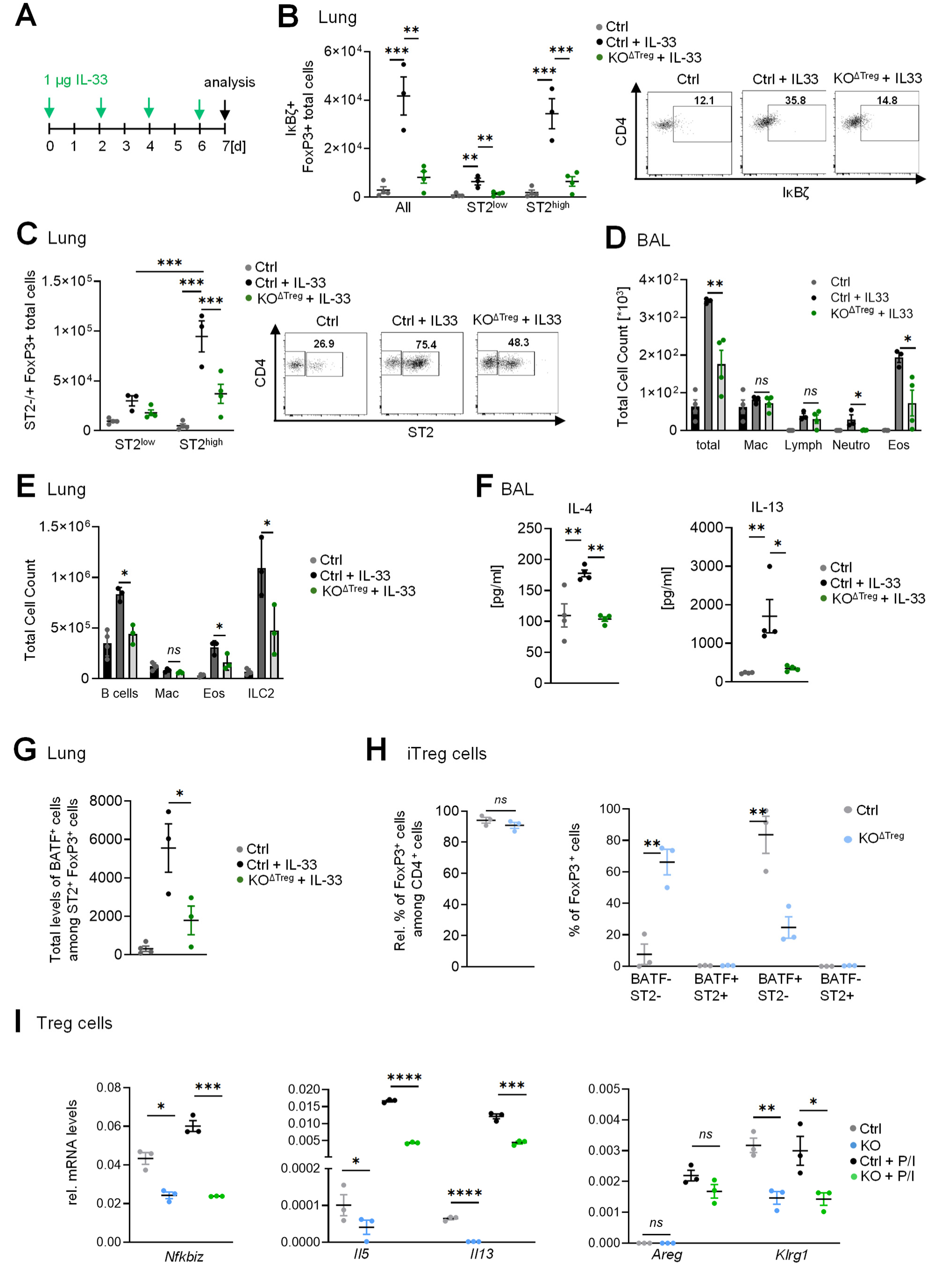
Treg-specific KO of IκBζ partially impairs IL-33-induced expansion of tissue-resident Treg cells and type 2 inflammation. **A.** IL-33 treatment scheme. Ctrl (*Nfkbiz* flox) and Treg-specific *Nfkbiz* KO mice (KO^ΔTreg^) received intraperitoneal injections of 1 µg IL–33 or a PBS control every second day. Mice were sacrificed and analysed on day 7. **B.** Flow cytometry analysis of IκBζ expression in lung Treg cells. Shown is the total number of IκBζ^+^ cells among all FoxP3^+^ Treg cells, subdivided into ST2^-^ and ST2^+^ Treg cells in the lungs of untreated Ctrl, and IL-33-treated Ctrl and KO^ΔTreg^ mice. N = 4 mice per group ± SEM. Right: representative dot plots of CD4 vs. IκBζ within FoxP3^+^ cells of all three experimental groups. **C.** Flow cytometry analysis of ST2 expression in lung Treg cells: Total number of ST2^-^ and ST2^+^ cells among FoxP3^+^ Treg cells in the lungs of untreated Ctrl, and IL-33-treated Ctrl and KO^ΔTreg^ mice. N = 4 mice per group ± SEM. Right: representative dot plots of CD4 *vs.* ST2 within FoxP3^+^ cells of all three experimental groups. **D.** Cellular composition of bronchoalveolar lavage (BAL). Shown are the total cell count (total) and numbers of macrophages (Mac), lymphocytes (Lymph), neutrophils (Neutro), and eosinophils (Eos) in untreated and IL-33-treated mice. N = 4 mice per group ± SEM. **E.** Flow cytometry analysis of lung-infiltrating immune cells. Shown is the total cell number of B cells, macrophages (Mac), eosinophils (Eos), and ILC2 cells in the lung tissue of untreated control, IL-33-treated control, and KO^ΔTreg^-treated mice. N = 4 mice per group ± SEM. **F.** Cytokine concentration of IL-4 and IL-13 in the BAL (bronchoalveolar lavage fluid). N = 4 mice per group ± SEM. **G.** Flow cytometry analysis of total cell numbers of BATF^+^ ST2^+^ Treg cells in the lung of untreated and treated mice. N = 4 mice per group ± SEM. **H.** Relative frequency of *in vitro* differentiated iTreg cells from splenic CD4^+^ T-cells of 11-16-week-old Ctrl and KO^ΔTreg^ mice, as well as BATF and ST2 protein expression in these iTreg cells, as assessed by flow cytometry. Shown is the mean of 3 biological replicates ± SEM per group. **I.** Gene expression analysis of MACS-bead-sorted Treg cells as in (H). Shown are relative mRNA levels in untreated and 2 h PMA/ionomycin (P/I)-stimulated Treg cells, normalized to *Actb*. N = 3 independent samples ± SEM. Significance was calculated using a one-way ANOVA with Tukeýs multiple comparison test. for the *in vivo* experiments, and a two-sided t-test for the in vitro experiments (\**p* ≤ 0.05, \*\**p* < 0.01, \*\*\**p* < 0.01, *ns* = not significant).

Previous reports showed that IL-33 not only promotes ST2^+^ Treg expansion, but also increases the abundance and activity of ILC2 and eosinophils, thereby contributing to type 2 inflammation ^35, 36^. Moreover, in wild–type mice, IL–33 has been demonstrated to reprogram ST2^+^ Treg cells towards a pro–inflammatory phenotype characterized by the production of Th2 cytokines ^33, 37^. Hence, we investigated the overall inflammatory response and especially type 2 inflammation in IL-33-treated control and *Nfkbiz* KO^ΔTreg^ mice. IL-33 treatment as well as the deletion of IκBζ in Treg cells caused only mild effects in the lymph nodes and spleen (Supplementary Figure S6C). However, we detected a diminished inflammation in the lungs of IL-33-treated *Nfkbiz* KO^ΔTreg^ mice compared to IL-33-treated controls, which was marked by a strong decline in total ILC2, neutrophil, B-cell, and eosinophil numbers in the bronchoalveolar lavage (BAL) (Figure 7D) and lung tissue (Figure 7E). Consistent with this observation, IL-33-treated *Nfkbiz* KO^ΔTreg^ mice displayed a global reduction in Th2-associated cytokine levels in the BAL (Figure 7F), accompanied by diminished Th2-driven goblet cell hyperplasia and reduced expression of *Il5* and *Il10* in the lung (Supplementary Figure S6D and S6E). Together, this data not only validates IL-33 as a trigger of IκBζ expression in ST2^+^Treg cells, but also proposes a model in which IκBζ triggers the conversion of Treg cells into disease-promoting Th2 cells that actively promote type 2 inflammation.

Since overexpression of IκBζ induces Th2 cytokine expression in Treg cells through modulation of BATF function, we next investigated whether the KO of IκBζ in Treg cells impairs their conversion into Th2-like effector cells by limiting BATF expression and function. Indeed, deletion of IκBζ in Treg cells significantly lowered the number of BATF^+^ cells within the ST2^+^ Treg cells upon IL-33 treatment (Figure 7G) as well as in *in vitro*-differentiated iTreg cells (Figure 7H). Furthermore, PMA/ionomycin treatment of wild-type magnetic bead-sorted Treg cells induced cell-intrinsic expression of *Il13*, *Il5*, and *Klgr1*, which was significantly impaired in *Nfkbiz* KO Treg cells (Figure 7I). Together, this implies that IL-33-mediated conversion of Treg cells into Th2-like Treg cells critically depends on IκBζ, most likely through its ability to modulate BATF function in Treg cells.

## Discussion

Treg cells are essential for maintaining peripheral immune tolerance and tissue homeostasis. Beyond their classical immunosuppressive functions, specialized subsets of tissue-resident Treg cells have emerged as critical regulators of tissue repair and local immune modulation^38^. While several transcriptional regulators governing tissue Treg cell differentiation have been identified, the mechanisms that stabilize or reprogram tissue Treg cells under inflammatory conditions remain incompletely understood ^5, 39^.

In this study, we identify the NF-κB transcriptional cofactor IκBζ as a central regulator of Treg cell plasticity. By combining Treg-specific gain- and loss-of-function approaches, we demonstrate that IκBζ acts as a molecular switch that converts Treg cells from immunosuppressive regulators into Th2-like effector cells. Constitutive expression of IκBζ in Treg cells triggered uncontrolled expansion of potentially functional impaired Treg cells, resulting in systemic Th2-dominated inflammation in barrier organs such as the lung, skin, and colon. Conversely, Treg-specific deletion of IκBζ restrained IL-33-driven Treg expansion and unexpectedly attenuated type 2 inflammation, particularly in the lung.

Mechanistically, our data establish BATF as the key transcriptional effector downstream of IκBζ in Treg cells. IκBζ overexpression increased BATF expression and chromatin occupancy at Th2-associated cytokine loci, including *Il4* and *Il13*, thereby inducing a cell-intrinsic Th2-like transcriptional program in Treg cells. This finding is particularly notable given that FoxP3, a repressor of Th2 differentiation in Treg cells ^8^, remained intact, indicating that IκBζ-driven reprogramming overrides canonical Treg lineage control mechanisms.

How IκBζ mechanistically enhances BATF function remains an important question. As IκBζ itself lacks any DNA-binding activity, it is thought to act as a bridging factor that recruits epigenetic modifiers to chromatin-associated transcription factor complexes. Previous work has shown that IκBζ can guide the BAF chromatin-remodeling complex to specific genomic loci, thereby increasing promoter accessibility, transcription factor binding, and ultimately facilitating gene expression ^17^. Intriguingly, BAF-mediated nucleosome remodeling has been identified as a prerequisite for BATF binding during acute activation of CD8⁺ effector T cells ^40^, and promotes IL-4 expression in conventional CD4⁺ T cells ^41^. Together with our data, these findings suggest a model in which IκBζ facilitates BATF-dependent transcription in Treg cells by promoting chromatin accessibility at Th2-associated gene loci. Interestingly, IκBζ did not strongly influence the differentiation of BATF-positive tissue Treg cells at steady-state, suggesting that this mechanism may be selectively engaged during inflammatory conditions, such as IL-33 signaling.

The earliest phenotypic changes in Treg-specific IκBζ-overexpressing mice comprised a Th2-like lung inflammation and an increased activity of B cells. BATF-expressing tissue-resident Treg cells have been implicated in tissue repair and resolution of inflammation ^5, 42^. Our data now demonstrate that in the presence of enforced IκBζ expression, tissue Treg cells acquire disease-promoting properties by producing Th2 cytokines and amplifying type 2 inflammation. This highlights a previously unappreciated role of IκBζ in the transition of tissue-resident Treg cells from protective regulators to inflammatory drivers, underscoring the importance of tightly controlled transcriptional programs in maintaining immune homeostasis.

Several reports have previously described a Treg-intrinsic expression of IL-4 and IL-13, for example, upon acute lung injury, intranasal application of IL-33, or in IL-33-stimulated visceral adipose tissue ^37, 43, 44^. Furthermore, BATF has previously been shown to induce IL-4 expression in T follicular helper cells ^30^, and IL-4 and IL-13 expression in Th2 cells upon helminth infection ^45^. Our results significantly expand these findings, as we show that IκBζ expression can trigger BATF-mediated induction of IL-4 and IL-13 in Treg cells. Consequently, this switch of Treg cells to Th2-like Treg cells provoked strong systemic Th2-like inflammation, marked by an increase in mucus-producing cells in the lung, fibrosis induction, and increased levels of IL-4 and IL-13 in various tissues, ultimately leading to B-cell class switching and production of large amounts of IgE. This finding implies that BATF-expressing Treg-like cells not only maintain tissue homeostasis by limiting tissue damage, but can also contribute to Th2-driven inflammation, depending on the co-expression of IκBζ.

Based on our findings and previous reports showing that IL-33 can convert Treg cells into Th2-like Treg cells ^33, 37^, we hypothesized that Treg-specific deletion of IκBζ preserves the activity of Treg cells, thereby limiting IL-33-induced type 2 inflammation. Indeed, while IL-33 robustly expanded ST2⁺ Treg cells in control mice as previously reported ^35^, Treg-specific deletion of IκBζ not only limits this expansion but also markedly reduced lung inflammation, eosinophilia, and ILC2 accumulation. This finding was unexpected, as ST2⁺ Treg cells are generally considered anti-inflammatory. Our data suggest that IL-33–induced IκBζ expression destabilizes Treg cell identity, enabling their conversion into Th2-like Treg cells that actively promote inflammation. In the absence of IκBζ, Treg cells appear to retain or even enhance their suppressive capacity, thereby improving the control of type 2 immune responses.

The precise mechanisms underlying the enhanced suppressive function of IκBζ-deficient Treg cells remain to be elucidated. One possibility is that IκBζ limits the expression of Treg-derived inhibitory molecules such as IL-10, TGF-β, ICOS, or OX40, which were previously shown to inhibit ILC2 function during acute lung injury ^36, 46^. Alternatively, Treg cells and ILC2 compete for IL-2 ^47^ and loss of IκBζ may increase IL-2 receptor expression or signaling in Treg cells, thereby limiting ILC2 expansion. Further studies are needed to elucidate the mechanism underlying the improved immunosuppressive function of IκBζ-deleted Treg cells.

From a translational perspective, our findings have important implications. Multiple therapeutic approaches, e.g., using retinoic acid or TNFR2 agonists, are being explored, aiming at an expansion of tissue-resident Treg cells for the treatment of graft-versus-host disease, wound healing defects, or fibrosis ^48–50^. Our findings highlight the importance of evaluating whether such interventions induce IκBζ expression, as this could inadvertently promote Treg destabilization and inflammation. In this context, TNFR2 agonists may be particularly attractive, as TNF signaling alone appears insufficient to induce IκBζ expression ^51^, potentially allowing expansion of tissue Treg cells without compromising their suppressive function.

Moreover, tissue-resident Treg cells accumulate in chronic inflammatory diseases such as asthma, atopic dermatitis, or inflammatory bowel disease ^52, 53^, yet their functional contribution to disease progression remains controversial. Our data suggest that enforced or sustained induction of IκBζ, potentially driven by inflammatory cytokines or tissue damage, may convert Treg cells from protective regulators into pathogenic effectors. Notably, aberrant IκBζ expression has already been implicated in diseases such as psoriasis ^19, 54^, an aggressive subtype of B-cell lymphoma ^55^, or melanoma ^56^, where it promotes chronic inflammation, disease progression, and therapy resistance.

In summary, our study identifies IκBζ as a critical determinant of Treg cell plasticity that controls BATF-dependent Th2-like reprogramming. Excessive IκBζ expression converts Treg cells into pathogenic effector cells, whereas its deletion preserves Treg stability and restrains inflammation under destabilizing conditions such as chronic IL-33 exposure. These findings uncover a previously unrecognized checkpoint in tissue Treg cell regulation and highlight IκBζ as a potential therapeutic target in type 2–driven inflammatory diseases.

## Material and Methods

### Mice

All mouse experiments were approved by the local animal ethics committee (Landesuntersuchungsamt Rheinland-Pfalz, G23-1-084). The following mouse lines were used: Ctrl: Control mice only possessing the doxycycline-inducible IκBζ overexpression cassette (called *Nfkbiz* Tet on, generated by InGenious, see Figure 1 and Supplementary Figure S1), Ctrl^ΔCre+^: Control mouse line only expressing the Sakaguchi FoxP3-Cre (Foxp3<tm1(cre)Saka>; provided by Ari Waisman, UMC Mainz), OE^ΔTreg^: Treg-specific IκBζ-overexpressing mice (homozygous for FoxP3-Cre and heterozygous or homozygous for *Nfkbiz* Teton). Control for Treg-specific *Nfkbiz* KO (Ctrl) is a mouse line harboring loxP sequences between the 4^th^ and the 7^th^ exon of the *Nfkbiz* gene (Nfkbiz<tm1.1Muta>, purchased from RIKEN Institute), KO^ΔTreg^ mice = Treg-specific IκBζ knockout mice, generated by crossing floxed *Nfkbiz* control mice with Bluestone FoxP3-Cre (Foxp3-EGFP/icre; provided by Caspar Ohnmacht, Helmholtz Center Munich). For IL-33 treatment, 8 - 12 week-old Ctrl and KO ^ΔTreg^ mice received intraperitoneal injections with 1 µg IL-33 (in 100 µL PBS, BioLegend, Cat. 580506) or PBS control every second day for 7 days. On day 7, mice were sacrificed and analyzed. In all experiments, female and male mice were used; the different experimental groups were always gender- and age-matched. Young mice were between 11 and 16 weeks old, and aged mice were older than 30 weeks.

### Isolation of Treg cells and doxycycline stimulation

Murine primary Treg cells were either obtained using the EasySep™ Mouse CD4+CD25+ Regulatory T Cell Isolation Kit II from Stemcell (Cat. 18783) or through FACS sorting. For FACS-based sorting, a single-cell suspension of splenocytes was first obtained as specified in the flow cytometry section. Next, T cells were pre-enriched using CD4 microbeads (Miltenyi Biotec, Cat. 130-117-043) and stained with the specific anti-mouse antibodies listed in Supplementary Table S1. For Dox treatment, Treg and T cells were isolated by magnetic bead isolation and incubated for 72 h in RPMI-1640 (Sigma-Aldrich, Cat. R8758-500ML) supplemented with 10% FCS (Sigma-Aldrich, Cat. F7524), antibiotics (Sigma-Aldrich, Cat. P0781), 50 µM β-mercaptoethanol (Thermo Fisher, Cat. 11528926), 1% non-essential amino acids (Thermo Fisher, Cat. 11140035), 5 ng/mL IL-2 (Immunotools, Cat. 12340024), and Dox (1 µg/mL, Biomol, Cat. Cay14422-1), at 5% CO_2_ and 37°C. Culture plates were pre-coated with 4 µg/mL anti-CD3 (BioLegend, Cat. 100340) and 2 µg/mL anti-CD28 (BioLegend, Cat. 102116).

### Generation of in vitro differentiated iTreg cells

Naïve CD4^+^ T cells were isolated from the spleen and lymph nodes using the Naïve CD4^+^ T Cell Isolation Kit from Stemcell (Cat. 19765). Cells were cultured in the same medium as mentioned above, without Dox addition. For the generation of iTreg cells, 1 µg/mL anti-IFNγ (Immunotools, Cat. 22853631), 1 µg/mL anti-IL-4 (BioLegend, Cat. 504102), and 15 ng/mL TGFꞵ (BioLegend, Cat. 763104) were added for 96 h. In some experiments, iTreg cells were additionally stimulated with P/I (50 ng/mL PMA and 500 ng/mL ionomycin) for 2 - 4 h, as indicated in the figure legends.

### In vitro suppression assay

CD4^+^CD25^-^ cells from wild-type mice and CD4^+^CD25^+^ cells from OE and Ctrl^ΔCre+^ mice were obtained with Stemcell Isolation Kits as described under “Isolation of Treg cells and doxycycline stimulation”. The proliferation of CD4^+^CD25^-^ cells was assessed by staining with the fluorescent dye CellTrace Violet (Invitrogen, Cat. C34557). Cells were stained at a density of 10^7^ cells/mL with 3 µM dye in PBS for 10 min at room temperature (RT) in the dark. The labeled cells were then seeded in anti-CD3 (4 µg/mL)-coated plates. Soluble anti-CD28 (2 µg/mL) was added at a density of 5 × 10^4 cells per well, and isolated Treg cells were added in various ratios (1:2, 1:4, or without Tregs). After 4 days of stimulation at 37°C and 5% CO₂, proliferation of the labeled CD4^+^CD25^-^ cells was assessed by flow cytometry.

### Histology

Tissue samples were fixed in 4% formaldehyde (Roth, Cat. P087.1) and embedded in paraffin, before 5-µm thick sections were prepared and stained with H&E as described ^57^. For the detection of mucus-producing goblet cells, sections were stained with periodic acid (Roth, Cat. 3257.1) and Schiff’s reagent (Avantor, Cat. 1.09033.0500), followed by incubation in SO_2_ water (6 mM) for 3 min. For collagen staining, sections were first stained with hematoxylin (Waldeck, Cat. 2E-032 and 2E052) and then incubated for 60 min in 0.1% sirius red (Waldeck, Cat. 1A-280) solution prepared in 1.2% aqueous picric acid (Waldeck, Cat. 3E-086; final pH: 2), followed by washing with deionized water and 1% acetic acid (Roth, Cat. 3738.4). Digitization was done with a NanoZoomer 2.0-HT scanner (Hamamatsu Photonics, Herrsching, Germany) using the software NDP.View2 (Hamamatsu Photonics). The degree of lung inflammation was assessed in a blind manner by an experienced observer. Five randomly selected areas were scored on a scale from 0 (no visible infiltrate around airway vessels and parenchyma) to 4 (multi-layered cellular infiltrates on nearly all visible vessels and airways). A detailed description of the scoring has been reported previously ^58^. Mucus-producing cells were quantified as the number of PAS-positive cells per millimeter of basal membrane on three representative airways per section.

### Immunoblot analysis of autoantibodies

Immunoblot analysis was carried out as described ^54^. To detect the presence of antibodies against endogenous antigens, protein lysates from organs of RAG2 knockout mice were prepared and separated by SDS-PAGE. After transfer to nitrocellulose membranes, membranes were incubated with 50 µL serum of Ctrl and OE ^ΔTreg^, diluted in PBS. Bound autoantibodies were detected using anti-mouse IgG coupled with horseradish peroxidase (Jackson ImmunoResearch, Cat. 115-035-003).

### Flow cytometry

Single-cell solutions containing 5 × 10^5 cells were used for flow cytometric staining in a 96-well plate (ThermoFisher, Cat. 277143). Unspecific binding was blocked by preincubation with 0.5 µL of Fc receptor-blocking antibodies (BioLegend, Cat. 101319) for 10 min at 4°C. Cells were then stained with the appropriate antibody mixtures for at least 15 min. Co-staining and gating strategy of the single cell types can be retrieved from Supplementary Table S1 and the respective Supplementary Figures. To identify dead cells, cells were incubated with eFluor780 fixable viability dye (ThermoFisher, Cat. 65-0865-14) during surface staining. For GC B cells, live and dead cells were discriminated using the dyes AmCyan (Ebioscience, Cat. 65-0866) and R780 (Ebioscience Cat. 65-0865) added for 10 min at 4°C, prior to surface staining. Intracellular epitopes were stained using the Foxp3/Transcription Factor Staining Buffer Set (ThermoFisher) according to the manual. To this end, cells were incubated with fixation/permeabilization buffer for 1 h at 4°C and then stained overnight with antibodies against intracellular epitopes in permeabilization buffer. For IκBζ staining, cells were fixed for 45 min at RT and incubated in Permbuffer containing 10% FCS for 10 min at RT. Cells were then resuspended in IκBζ antibody mix for 45 min at RT. After staining, cells were washed and incubated in FACS buffer until measurement. At the beginning of each analysis, doublets and debris were excluded based on FSC-A/FSC-H, followed by exclusion of dead cells and gating on CD45⁺ leukocytes. All analyses were performed using FlowJo v10.10.0 software with representative gating shown in the respective Supplementary Figures.

### Gene expression analysis

Gene expression analysis was performed as described, using custom-designed primers (Supplementary Table S2) ^56^. For relative mRNA quantification, threshold cycle (Ct) values of target genes were normalized to the Ct-values of the reference gene *Actb* or *Hprt1* using the 2^(-ΔCt)^ method.

#### ELISA

Quantitative levels of IL-4 and IL-13 were determined in lung homogenates, BAL, and serum samples using the ELISA MAX^TM^ Deluxe Set Mouse IL-4 (BioLegend, Cat. 431102) and Mouse IL-13 DuoSet ELISA (R&D Systems, Cat. DY413) according to the manufacturer’s instructions. Quantitative levels of murine IgG1 and IgE were determined in serum samples using Immuno 96-well plates (Thermo Fisher, Cat. 442404) coated with goat anti-mouse IgG1 (SouthernBiotech, Cat. 1070-01) or rat anti-mouse IgE (CliniSciences, Cat. 1130-01). After washing, self-made standard and serum samples were added for 1 hour at 37°C. Plates were washed again and incubated for 1 hour at 37°C with the detection antibodies rat anti-mouse IgE-BIOT (SouthernBiotech, Cat. 1144-08) or rat anti-mouse IgE-BIOT (CliniSciences, Cat. 1130-08), followed by incubation with streptavidin antibody (Sigma, Cat. 11089161001) for 30-60 min at RT. Lastly, the biotinylated substrate (Santa Cruz, Cat. 206099) was added, and the reaction was stopped once the highest standard reached an OD of 2.5. Absorbance was measured using a microplate reader (Hidex Sense, Hidex), and cytokine concentrations were calculated by fitting standard curves using a four-parameter logistic regression.

#### Bulk RNA sequencing

Total RNA library preparation and sequencing were carried out by Novogene (Munich, Germany). Following library quantification, sequencing was performed on the Illumina NovaSeq X Plus system using pair-end 150 bp reads (PE150), with sequencing depth adjusted according to effective library concentration and required data output. Raw sequencing reads were processed using CLC Genomics Workbench (v25,0.1; Qiagen). Read alignment to the mouse reference genome (GRCm39, assembly GRCm39.111) was conducted by the NGS core facility at the Research Centre for Immunology using default CLC parameters (mismatch cost = 2; insertion cost = 3; deletion cost = 3; length fraction = 0.8; similarity fraction = 0.8; global alignment disabled; strand-specific = both; library type = bulk; maximum hits per read = 10; paired reads counted as two = no; broken pairs ignored). Differential gene expression was defined by an absolute fold change > 2, an absolute difference exceeding 4, and an adjusted *P* value ≤ 0.05, when comparing WT and IκBζ overexpression samples using the proportion-based Baggerleýs test. Raw and processed sequencing data have been deposited in the GEO database under accession number GSE318157. Heatmaps were generated using the Morpheus software platform (Broad Institute) (https://software.broadinstitute.org/morpheus).

#### Single-cell RNA sequencing

Spleens were mashed through a 70 µm strainer to obtain single-cell suspensions. To prevent nonspecific antibody binding, Fc blocking reagent (Miltenyi Biotec, Cat. 130-092-575) was added. CD45^+^ cells were subsequently isolated by MACS bead sorting, according to the manual (Miltenyi Biotec, Cat. 130-052-301). Afterwards, the enriched CD45^+^ cells were labeled with TotalSeqC anti-mouse Hashtagging antibodies (Biolegend, Cat. 155861, 155863) for 25 min at 4°C. To enhance labeling efficiency, cells were washed 3–5 times with 500 μL FACS buffer after staining. To prevent aggregation during staining, the antibody mix was centrifuged at 14,000g for 10 min at 4°C, and the supernatant was transferred to a new tube. Labeled target cells were transferred into a single 1.5 mL tube containing 350 μL PBS + 0.05% BSA. After washing and supplementation with master mix and beads, samples were loaded onto a 10X Chromium Next GEM Chip K (10X Genomics, Cat. PN-1000286) and processed using a 10X Chromium Controller. Subsequently, cDNA amplification was performed using the Chromium Next GEM Single Cell 5’ Reagent Kit v2 (10X Genomics, Cat. PN-1000263) and the 5’ Feature Barcode Kit. Libraries for GEX and CSP were prepared according to the Library Construction Protocol (10X Genomics, Cat. PN-1000190). The fragment length composition was evaluated via electrophoretic separation of the samples using Agilent High Sensitivity D1000 ScreenTape assay and reagents (2200 Tapestation Controller, Agilent).

The generated scRNA libraries were sequenced by Novogene on an Illumina NextSeq 500/550 platform with a 150-cycle high-output cartridge. Sequencing was performed according to the manufacturer’s protocol: paired-end, with 26 base pairs (bp) for Read1, 90 bp for Read2, and 10 bp for the Index i5 and Index i7 (PE 26-10-10-90). After sequencing, the FASTQ files were aligned to the reference genome refdata-gex-mm10-2020-A using CellRanger (7.1.0), a software tool designed for single-cell sequencing datasets generated with 10X Genomics chemistry. Integration of GEX and Feature Barcode information was performed using CellRanger Multi, a method developed for processing scRNA samples with specific multiplexing antibodies. This approach supports the analysis of 3’ multiplexed data. Since the 3’ and 5’ assays capture different transcript ends, and the 5’ chemistry was employed to generate the GEX and CSP libraries, modifications to the CellRanger Multi pipeline were made to ensure dataset compatibility. The pipeline was executed to assign cells to individual samples by combining the GEX and CSP libraries. Output files were used for further analysis in R. The sequencing raw data and called peaks can be found under the GEO accession number GSE318157.

#### Chromatin immunoprecipitation and sequencing

Chromatin immunoprecipitation (ChIP) assays were performed as previously described ^57^. Briefly, chromatin was crosslinked with 0.25 M di-(N-succinimidyl) glutarate (Thermo Fisher, Cat. 20593) for 45 min at RT, followed by crosslinking with 1% formaldehyde (Thermo Fisher, Cat. 28906). After sonication, chromatin was incubated overnight at 4°C on a rotator with protein G-coupled Dynabeads (Thermo Fisher, Cat. 10003D) and anti-IκBζ (Cell Signaling, Cat. 76041), anti-BATF (Abcam, Cat. ab236876), or IgG control antibody (Abcam, ab46540). For validation of the ChIP by qPCR, self-designed primers mapping to BATF-binding sites were used, according to publicly available BATF ChIP sequencing data sets, retrieved from Cistrome Finder (Supplementary Table S3). Quantitative PCR was performed using Maxima SYBR Green Master Mix (Thermo Fisher, Cat. K0221), and ChIP enrichment was calculated as a percentage of input as previously described.

For ChIP sequencing, sequencing libraries were prepared by Novogene using the NEBNext® Ultra™ II DNA Library Prep Kit for Illumina®, according to the manual (without size selection and 15 cycles of final PCR). All quality controls were done using Invitrogen’s Qubit HS assay, and library size was determined using Agilent’s TapeStation 4150. Barcoded libraries were sequenced on an Illumina NovaSeq X Plus (150 Cycles; paired-end) at Novogene (Munich, Germany). Computational analysis was performed using the nf-core chipseq pipeline (v 2.0.0, https://nf-co.re/chipseq/2.0.0/). The pipeline contains an automated workflow integrating state-of-the-art tools. Initial quality control and adapter trimming of FASTQ files were performed by FastQC (https://www.bioinformatics.babraham.ac.uk/projects/fastqc/) and Trim Galore (https://github.com/FelixKrueger/TrimGalore), respectively. Bowtie2 (https://bowtie-bio.sourceforge.net/bowtie2/index.shtml) was selected for read alignment to the reference genome assembly (mm10) from the available aligners in the pipeline. Downstream manipulation and QC were performed on generated BAM files using Picard (https://broadinstitute.github.io/picard/), SAMtools (https://www.htslib.org/), BAMtools (https://github.com/pezmaster31/bamtools), and Preseq (https://smithlabresearch.org/software/preseq/). BEDtools (https://bedtools.readthedocs.io/en/latest/) was utilized to remove the blacklisted regions of the genome. Narrow peaks calling was performed using MACS2 (https://github.com/macs3-project/MACS/) with read length set to 150. The narrowPeak bed files were also converted to BigWig format for visualization via IGV using bedGraphToBigWig (https://www.encodeproject.org/software/bedgraphtobigwig/).To identify peaks in the promoter regions of the genome, we utilized the refTSS database (https://reftss.riken.jp/). Peaks 5 kb up- and downstream of the transcription start site (tss) were selected for subsequent analysis. Target prediction was done using the BETA-minus software from Cistrome (cistrome.org). The sequencing raw data and called peaks can be found under the GEO accession number: GSE318156.

#### Statistics

All experiments were performed with at least three independent biological replicates. Statistical significance was measured using a two-tailed Student’s *t*-test, with p ≤ 0.05 being considered statistically significant (* p < 0.05; ** p < 0.01; *** p < 0.001 and **** p < 0.0001), if not otherwise indicated in the figure legends. Data from *in vitro* experiments are presented as mean ± standard deviation (SD), while data obtained from *in vivo* experiments are shown as the mean ± standard error (SEM). All data plots were generated using GraphPad Prism.

## Acknowledgments

The study was supported by grants from the Peter Hans Hofschneider professorship of Molecular Medicine, funded by the Foundation for Experimental Biomedicine (D.K.), and the Deutsche Forschungsgemeinschaft TRR355/1 (project number 490846870, A01, A05, A06, A08) (to M.D., N.H., V.H., A.W., and D.K.), TRR156/3 (project number 246807620, B09, C08; to D.K. and A.W.), and SFB 1292/3 (project number 318346496, TP19 and TP08 (M.D. and D.K). D.K. is further supported by the Rise up! program of the Boehringer Ingelheim Foundation (BIS) and the ReALity initiative of the state of Rhineland-Palatinate. We thank Claudia Braun from the Histology Core Facility, Bonny Adami from the Biobank (University Medical Center of the Johannes Gutenberg-University Mainz), Anna Stastny, and Elena Zurkowski (University Medical Center of the Johannes Gutenberg-University Mainz) for technical assistance.

## Author contributions

T.K., A.-M. K.M., A.K., E.S. Z.E., D.-M. M., S.-S. H., S.R., N.B., and M.K. performed experiments and data analysis. M.D. and K.S. performed single-cell sequencing and analyzed the data. N.H. performed B-cell assays. A.W. helped to design the analysis of T cells. S.R. designed and analyzed experiments in the lung. S.H., K.S-O., and D.K. designed the IκBζ overexpression mouse model. D.K. designed the study and is the lead investigator for all animal experiments. S.R. and D.K. drafted the first version of the manuscript. All authors reviewed the manuscript and provided feedback.

## References

1. Dikiy S, Rudensky AY. Principles of regulatory T cell function. Immunity. Feb 14 2023;56(2):240–255. doi:10.1016/j.immuni.2023.01.004

2. Munoz-Rojas AR, Mathis D. Tissue regulatory T cells: regulatory chameleons. Nat Rev Immunol. Sep 2021;21(9):597–611. doi:10.1038/s41577-021-00519-w

3. Jugder BE, Park E, Du L, et al. Tissue-specific roles of regulatory T cells: mechanisms of suppression and beyond along with emerging therapeutic insights in autoimmune indications. Front Immunol. 2025;16:1650451. doi:10.3389/fimmu.2025.1650451

4. Raheem A, Khan I, Ahmad I, et al. The emerging role of tissue regulatory T cells in tissue repair and regeneration. Front Immunol. 2025;16:1640113. doi:10.3389/fimmu.2025.1640113

5. Delacher M, Imbusch CD, Hotz-Wagenblatt A, et al. Precursors for Nonlymphoid-Tissue Treg Cells Reside in Secondary Lymphoid Organs and Are Programmed by the Transcription Factor BATF. Immunity. Feb 18 2020;52(2):295–312 e11. doi:10.1016/j.immuni.2019.12.002

6. Delacher M, Schmidl C, Herzig Y, et al. Rbpj expression in regulatory T cells is critical for restraining T(H)2 responses. Nat Commun. Apr 8 2019;10(1):1621. doi:10.1038/s41467-019-09276-w

7. Griesenauer B, Paczesny S. The ST2/IL-33 Axis in Immune Cells during Inflammatory Diseases. Front Immunol. 2017;8:475. doi:10.3389/fimmu.2017.00475

8. Van Gool F, Nguyen MLT, Mumbach MR, et al. A Mutation in the Transcription Factor Foxp3 Drives T Helper 2 Effector Function in Regulatory T Cells. Immunity. Feb 19 2019;50(2):362–377 e6. doi:10.1016/j.immuni.2018.12.016

9. Heath VL, Murphy EE, Crain C, Tomlinson MG, O’Garra A. TGF-beta1 down-regulates Th2 development and results in decreased IL-4-induced STAT6 activation and GATA-3 expression. Eur J Immunol. Sep 2000;30(9):2639–49. doi:10.1002/1521-4141(200009)30:9<2639::AID-IMMU2639>3.0.CO;2-7

10. Mantel PY, Kuipers H, Boyman O, et al. GATA3-driven Th2 responses inhibit TGF-beta1-induced FOXP3 expression and the formation of regulatory T cells. PLoS Biol. Dec 2007;5(12):e329. doi:10.1371/journal.pbio.0050329

11. Annemann M, Plaza-Sirvent C, Schuster M, et al. Atypical IkappaB proteins in immune cell differentiation and function. Immunol Lett. Mar 2016;171:26–35. doi:10.1016/j.imlet.2016.01.006

12. Okamoto K, Iwai Y, Oh-hora M, et al. I kappa B zeta regulates T(H)17 development by cooperating with ROR nuclear receptors. Nature. Apr 29 2010;464(7293):1381-U13. doi:10.1038/nature08922

13. Miyake T, Satoh T, Kato H, et al. IkappaBzeta is essential for natural killer cell activation in response to IL-12 and IL-18. Proc Natl Acad Sci U S A. Oct 12 2010;107(41):17680–5. doi:10.1073/pnas.1012977107

14. Horber S, Hildebrand DG, Lieb WS, et al. The Atypical Inhibitor of NF-kappaB, IkappaBzeta, Controls Macrophage Interleukin-10 Expression. J Biol Chem. Jun 10 2016;291(24):12851–12861. doi:10.1074/jbc.M116.718825

15. Lorscheid S, Müller A, Löffler J, et al. Keratinocyte-derived IkappaBzeta drives psoriasis and associated systemic inflammation. JCI Insight. Nov 14 2019;4(22)doi:10.1172/jci.insight.130835

16. Yamamoto M, Yamazaki S, Uematsu S, et al. Regulation of Toll/IL-1-receptor-mediated gene expression by the inducible nuclear protein IkappaBzeta. Nature. Jul 8 2004;430(6996):218–22. doi:10.1038/nature02738

17. Tartey S, Matsushita K, Vandenbon A, et al. Akirin2 is critical for inducing inflammatory genes by bridging I kappa B-zeta and the SWI/SNF complex. Embo J. Oct 16 2014;33(20):2332–2348. doi:10.15252/embj.201488447

18. Yamazaki S. The Nuclear NF-kappaB Regulator IkappaBzeta: Updates on Its Molecular Functions and Pathophysiological Roles. Cells. Aug 31 2024;13(17)doi:10.3390/cells13171467

19. Johansen C, Mose M, Ommen P, et al. I kappa B zeta is a key driver in the development of psoriasis. P Natl Acad Sci USA. Oct 27 2015;112(43):E5825–E5833. doi:10.1073/pnas.1509971112

20. Yamazaki S, Inohara N, Ohmuraya M, et al. IkappaBzeta controls IL-17-triggered gene expression program in intestinal epithelial cells that restricts colonization of SFB and prevents Th17-associated pathologies. Mucosal Immunol. Jun 2022;15(6):1321–1337. doi:10.1038/s41385-022-00554-3

21. MaruYama T. TGF-beta-induced IkappaB-zeta controls Foxp3 gene expression. Biochem Biophys Res Commun. Aug 21 2015;464(2):586–9. doi:10.1016/j.bbrc.2015.07.013

22. MaruYama T, Kobayashi S, Ogasawara K, Yoshimura A, Chen W, Muta T. Control of IFN-gamma production and regulatory function by the inducible nuclear protein IkappaB-zeta in T cells. J Leukoc Biol. Sep 2015;98(3):385–93. doi:10.1189/jlb.2A0814-384R

23. Schiering C, Krausgruber T, Chomka A, et al. The alarmin IL-33 promotes regulatory T-cell function in the intestine. Nature. Sep 25 2014;513(7519):564–568. doi:10.1038/nature13577

24. Vasanthakumar A, Moro K, Xin A, et al. The transcriptional regulators IRF4, BATF and IL-33 orchestrate development and maintenance of adipose tissue-resident regulatory T cells. Nat Immunol. Mar 2015;16(3):276–85. doi:10.1038/ni.3085

25. Ohto-Ozaki H, Hayakawa M, Kamoshita N, Maruyama T, Tominaga SI, Ohmori T. Induction of IkappaBzeta Augments Cytokine and Chemokine Production by IL-33 in Mast Cells. J Immunol. Apr 15 2020;204(8):2033–2042. doi:10.4049/jimmunol.1900315

26. Sahnoon L, Bajbouj K, Mahboub B, Hamoudi R, Hamid Q. Targeting IL-13 and IL-4 in Asthma: Therapeutic Implications on Airway Remodeling in Severe Asthma. Clin Rev Allergy Immunol. Apr 21 2025;68(1):44. doi:10.1007/s12016-025-09045-2

27. Pene J, Rousset F, Briere F, et al. IgE production by normal human lymphocytes is induced by interleukin 4 and suppressed by interferons gamma and alpha and prostaglandin E2. Proc Natl Acad Sci U S A. Sep 1988;85(18):6880–4. doi:10.1073/pnas.85.18.6880

28. Spath S, Roan F, Presnell SR, Hollbacher B, Ziegler SF. Profiling of Tregs across tissues reveals plasticity in ST2 expression and hierarchies in tissue-specific phenotypes. iScience. Sep 16 2022;25(9):104998. doi:10.1016/j.isci.2022.104998

29. Chen EY, Tan CM, Kou Y, et al. Enrichr: interactive and collaborative HTML5 gene list enrichment analysis tool. BMC Bioinformatics. Apr 15 2013;14:128. doi:10.1186/1471-2105-14-128

30. Sahoo A, Alekseev A, Tanaka K, et al. Batf is important for IL-4 expression in T follicular helper cells. Nat Commun. Aug 17 2015;6:7997. doi:10.1038/ncomms8997

31. Daly AE, Yeh G, Soltero S, Smale ST. Selective regulation of a defined subset of inflammatory and immunoregulatory genes by an NF-kappaB p50-IkappaBzeta pathway. Genes Dev. Jun 25 2024;doi:10.1101/gad.351630.124

32. Zhou X, Jeker LT, Fife BT, et al. Selective miRNA disruption in T reg cells leads to uncontrolled autoimmunity. J Exp Med. Sep 1 2008;205(9):1983–91. doi:10.1084/jem.20080707

33. Chen CC, Kobayashi T, Iijima K, Hsu FC, Kita H. IL-33 dysregulates regulatory T cells and impairs established immunologic tolerance in the lungs. J Allergy Clin Immunol. Nov 2017;140(5):1351–1363 e7. doi:10.1016/j.jaci.2017.01.015

34. Matta BM, Reichenbach DK, Zhang X, et al. Peri-alloHCT IL-33 administration expands recipient T-regulatory cells that protect mice against acute GVHD. Blood. Jul 21 2016;128(3):427–39. doi:10.1182/blood-2015-12-684142

35. Ngo Thi Phuong N, Palmieri V, Adamczyk A, et al. IL-33 Drives Expansion of Type 2 Innate Lymphoid Cells and Regulatory T Cells and Protects Mice From Severe, Acute Colitis. Front Immunol. 2021;12:669787. doi:10.3389/fimmu.2021.669787

36. Stockis J, Yip T, Moreno-Vicente J, et al. Cross-talk between ILC2 and Gata3(high) T(regs) locally constrains adaptive type 2 immunity. Sci Immunol. Jul 19 2024;9(97):eadl1903. doi:10.1126/sciimmunol.adl1903

37. Siede J, Frohlich A, Datsi A, et al. IL-33 Receptor-Expressing Regulatory T Cells Are Highly Activated, Th2 Biased and Suppress CD4 T Cell Proliferation through IL-10 and TGFbeta Release. Plos One. 2016;11(8):e0161507. doi:10.1371/journal.pone.0161507

38. Loffredo LF, Savage TM, Ringham OR, Arpaia N. Treg-tissue cell interactions in repair and regeneration. J Exp Med. Jun 3 2024;221(6)doi:10.1084/jem.20231244

39. Hayatsu N, Miyao T, Tachibana M, et al. Analyses of a Mutant Foxp3 Allele Reveal BATF as a Critical Transcription Factor in the Differentiation and Accumulation of Tissue Regulatory T Cells. Immunity. Aug 15 2017;47(2):268–283 e9. doi:10.1016/j.immuni.2017.07.008

40. McDonald B, Chick BY, Ahmed NS, et al. Canonical BAF complex activity shapes the enhancer landscape that licenses CD8(+) T cell effector and memory fates. Immunity. Jun 13 2023;56(6):1303–1319 e5. doi:10.1016/j.immuni.2023.05.005

41. Jeong SM, Lee C, Lee SK, Kim J, Seong RH. The SWI/SNF chromatin-remodeling complex modulates peripheral T cell activation and proliferation by controlling AP-1 expression. J Biol Chem. Jan 22 2010;285(4):2340–50. doi:10.1074/jbc.M109.026997

42. Delacher M, Simon M, Sanderink L, et al. Single-cell chromatin accessibility landscape identifies tissue repair program in human regulatory T cells. Immunity. Apr 13 2021;54(4):702–720 e17. doi:10.1016/j.immuni.2021.03.007

43. Liu Q, Dwyer GK, Zhao Y, et al. IL-33-mediated IL-13 secretion by ST2+ Tregs controls inflammation after lung injury. JCI Insight. Mar 21 2019;4(6)doi:10.1172/jci.insight.123919

44. Hemmers S, Schizas M, Rudensky AY. T reg cell-intrinsic requirements for ST2 signaling in health and neuroinflammation. J Exp Med. Feb 1 2021;218(2)doi:10.1084/jem.20201234

45. Bao K, Carr T, Wu J, et al. BATF Modulates the Th2 Locus Control Region and Regulates CD4+ T Cell Fate during Antihelminth Immunity. J Immunol. Dec 1 2016;197(11):4371–4381. doi:10.4049/jimmunol.1601371

46. Rigas D, Lewis G, Aron JL, et al. Type 2 innate lymphoid cell suppression by regulatory T cells attenuates airway hyperreactivity and requires inducible T-cell costimulator-inducible T-cell costimulator ligand interaction. J Allergy Clin Immunol. May 2017;139(5):1468–1477 e2. doi:10.1016/j.jaci.2016.08.034

47. Molofsky AB, Van Gool F, Liang HE, et al. Interleukin-33 and Interferon-gamma Counter-Regulate Group 2 Innate Lymphoid Cell Activation during Immune Perturbation. Immunity. Jul 21 2015;43(1):161–74. doi:10.1016/j.immuni.2015.05.019

48. Torrey H, Kuhtreiber WM, Okubo Y, et al. A novel TNFR2 agonist antibody expands highly potent regulatory T cells. Sci Signal. Dec 8 2020;13(661)doi:10.1126/scisignal.aba9600

49. Chopra M, Biehl M, Steinfatt T, et al. Exogenous TNFR2 activation protects from acute GvHD via host T reg cell expansion. J Exp Med. Aug 22 2016;213(9):1881–900. doi:10.1084/jem.20151563

50. Steele H, Cheng J, Willicut A, et al. TNF superfamily control of tissue remodeling and fibrosis. Front Immunol. 2023;14:1219907. doi:10.3389/fimmu.2023.1219907

51. Kubelbeck T, Wichmann NO, Raj T, et al. Regulation and Function of the Atypical IkappaBs-Bcl-3, IkappaB(NS), and IkappaBzeta-in Lymphocytes and Autoimmunity. Eur J Immunol. May 2025;55(5):e202451273. doi:10.1002/eji.202451273

52. Roesner LM, Floess S, Witte T, Olek S, Huehn J, Werfel T. Foxp3(+) regulatory T cells are expanded in severe atopic dermatitis patients. Allergy. Dec 2015;70(12):1656–60. doi:10.1111/all.12712

53. Kryczek I, Wu K, Zhao E, et al. IL-17+ regulatory T cells in the microenvironments of chronic inflammation and cancer. J Immunol. Apr 1 2011;186(7):4388–95. doi:10.4049/jimmunol.1003251

54. Müller A, Hennig A, Lorscheid S, et al. IkappaBzeta is a key transcriptional regulator of IL-36-driven psoriasis-related gene expression in keratinocytes. Proc Natl Acad Sci U S A. Sep 17 2018;doi:10.1073/pnas.1801377115

55. Nogai H, Wenzel SS, Hailfinger S, et al. IkappaB-zeta controls the constitutive NF-kappaB target gene network and survival of ABC DLBCL. Blood. Sep 26 2013;122(13):2242–50. doi:10.1182/blood-2013-06-508028

56. Kolb A, Kulis-Mandic AM, Klein M, et al. Constitutive expression of the transcriptional co-activator IkappaBzeta promotes melanoma growth and immunotherapy resistance. Nat Commun. Jun 25 2025;16(1):5387. doi:10.1038/s41467-025-60929-5

57. Müller A, Dickmanns A, Resch C, et al. The CDK4/6-EZH2 pathway is a potential therapeutic target for psoriasis. J Clin Invest. Nov 2 2020;130(11):5765–5781. doi:10.1172/JCI134217

58. Reuter S, Lemmermann NAW, Maxeiner J, et al. Coincident airway exposure to low-potency allergen and cytomegalovirus sensitizes for allergic airway disease by viral activation of migratory dendritic cells. PLoS Pathog. Mar 2019;15(3):e1007595. doi:10.1371/journal.ppat.1007595

59. Evangelista JE, Xie Z, Marino GB, Nguyen N, Clarke DJB, Ma’ayan A. Enrichr-KG: bridging enrichment analysis across multiple libraries. Nucleic Acids Res. Jul 5 2023;51(W1):W168–W179. doi:10.1093/nar/gkad393

