## Supplementary Files for "Treg-derived IκBζ promotes their conversion into Th2-like effectors and drives type 2 inflammation via BATF"

### Supplementary information

**Supplementary Figure S1.** Detailed description of the cloning cassette for the generation of $OE^{\Delta Treg}$ mice.

**Supplementary Figure S2.** Additional data on Treg cell frequencies in aged mice and on FoxP3 expression in $OE^{\Delta Treg}$ mice.

**Supplementary Figure S3.** Additional data on Th2-associated disease pattern in $OE^{\Delta Treg}$ mice.

**Supplementary Figure S4.** Additional single-cell sequencing data of spleens from young and aged Ctrl and $OE^{\Delta Treg}$ mice.

**Supplementary Figure S5.** Supporting data on the modulation of BATF by IκBζ in Treg cells.

**Supplementary Figure S6.** Extended characterization of *Nfkbiz* KO mice ($KO^{\Delta Treg}$) with and without IL-33 treatment.

**Supplementary Table S1.** Flow cytometry antibodies.

**Supplementary Table S2.** Gene expression primer.

**Supplementary Table S3.** ChIP primer.

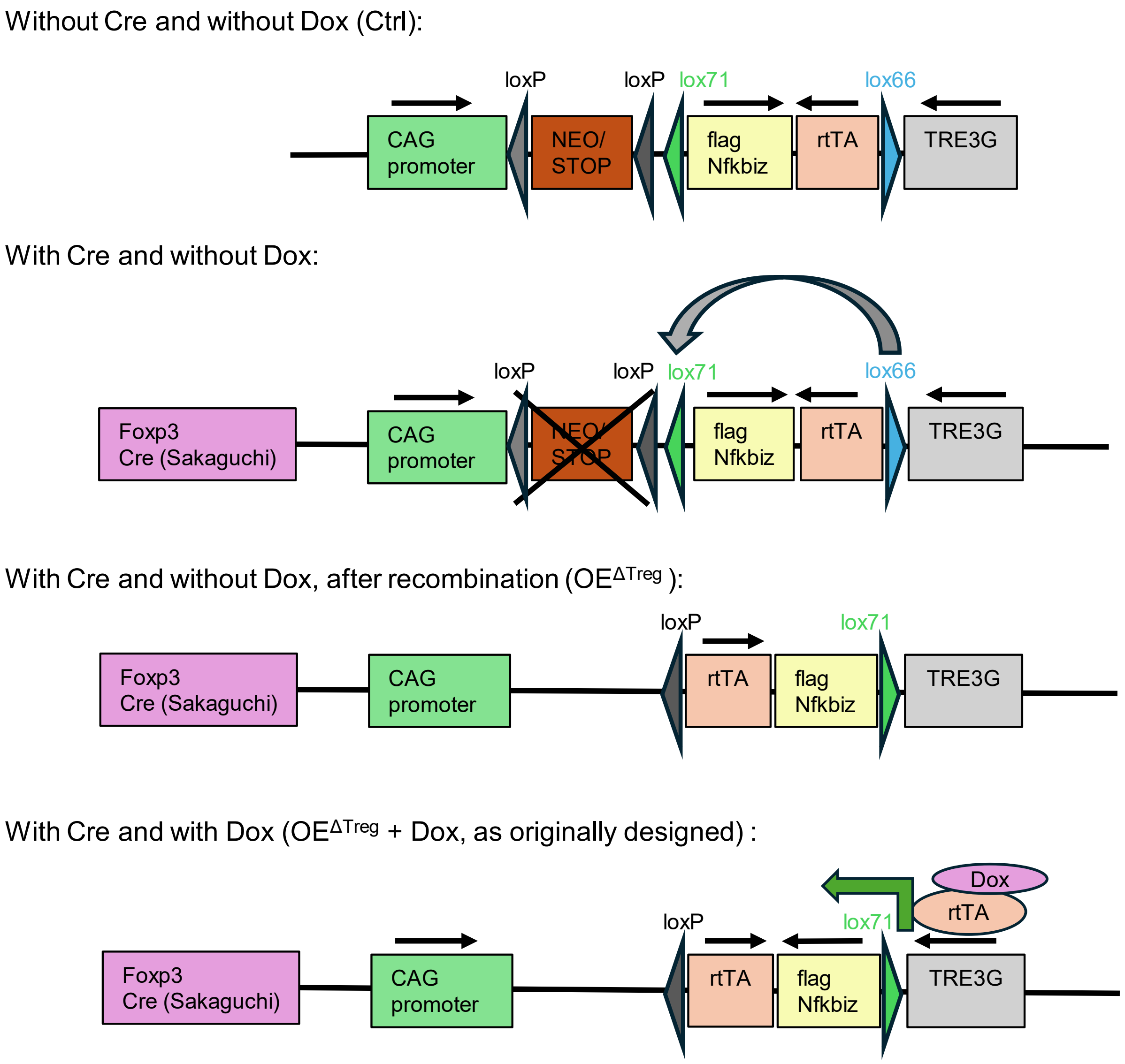


**Supplementary Figure S1. Detailed description of the cloning cassette for the generation of OE$^{\Delta Treg}$ mice.** The overexpression cassette is composed of a CAG promoter, a Neomycin Stop cassette flanked by loxP sites (NEO/STOP), cDNA encoding flag-tagged mouse *Nfkbiz* (NM_001159394.1) (flag-*Nfkbiz*), a reverse tetracycline-controlled transactivator (rtTA), and a doxycycline-inducible promoter (TRE3G). Importantly, the flag-*Nfkbiz* and rtTA cassette is flanked by lox71 and lox66 sites, enabling a Cre recombinase inversion of the cassette in the presence of a Cre Recombinase, in this case under the control of a Foxp3-promoter element. Due to the inversion of the cassette, rtTA is expected to be (Treg-specifically) constitutively expressed, whereas flag-*Nfkbiz* should be under the control of the doxycyclin inducible promoter. However, as Treg-specific overexpression of *Nfkbiz* was observed in the absence of doxycycline, these findings indicate that the intended cassette inversion using lox71/lox66 did not occur as designed.

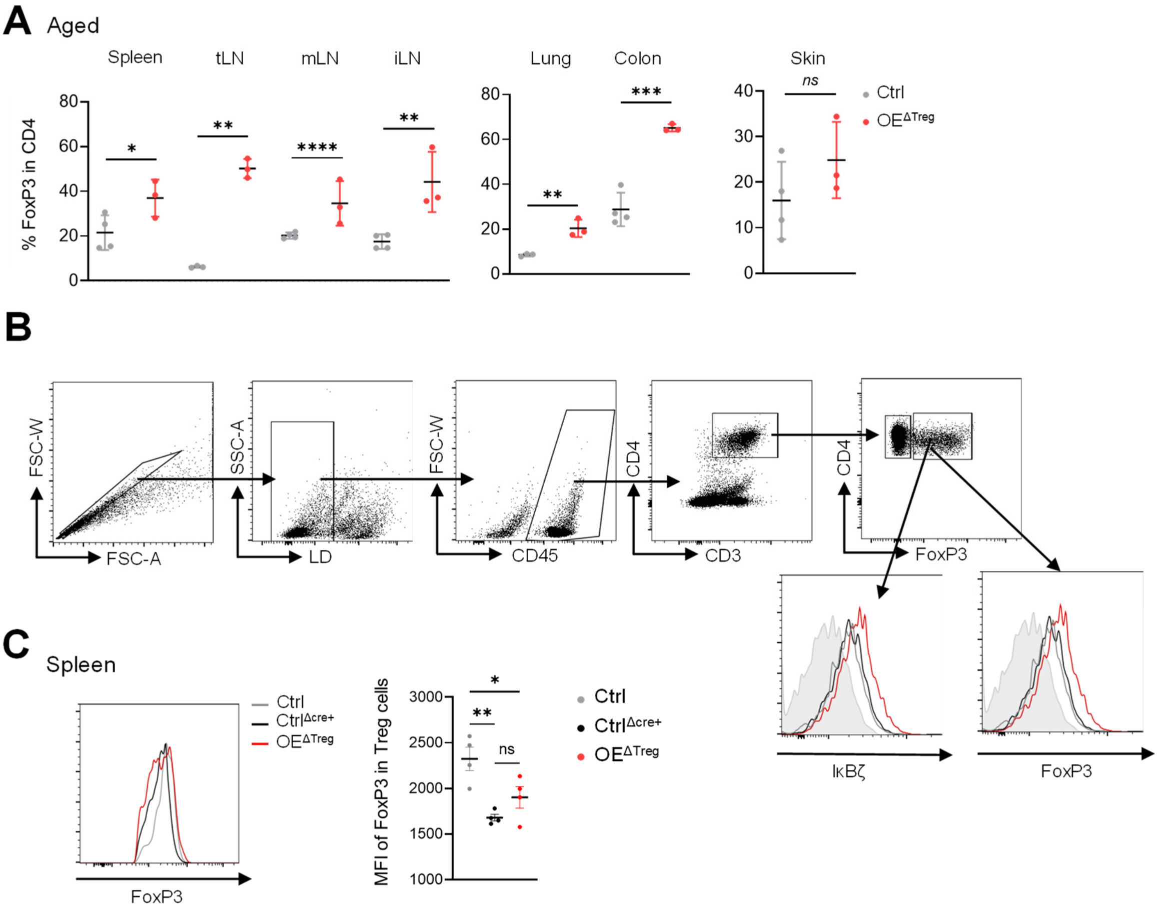


**Supplementary Figure S2. Additional data on Treg cell frequencies in aged mice and on FoxP3 expression in OE$^{\Delta Treg}$ mice. A.** Relative amount of FoxP3$^+$ Treg cells in various tissues of aged mice (over 30 weeks). FACS analysis of Treg cells from spleen and lymph nodes (tLN = tracheal lymph node, mLN = mesenteric lymph node, iLN = inguinal lymph node), lung, colon, and skin, shown as the percentage of FoxP3$^+$ cells among all CD4$^+$ cells, after pre-gating on viable cells. N = 3 – 4 mice per group ± SEM. **B.** Gating strategy to analyze Treg cells: Following exclusion of doublets and dead cells, CD45$^+$ leukocytes were selected. Within this population, T helper cells were identified as CD3$^+$ and CD4$^+$. Treg cells were characterized as FoxP3$^+$ CD3$^+$ CD4$^+$ cells. To assess the expression of IκBζ and FoxP3 in Treg cells, the MFI was further analyzed. **C.** Data from the spleen of Ctrl, Ctrl$^{\Delta cre+}$, and OE$^{\Delta Treg}$ mice. *Left:* Example plot of the FoxP3 staining normalized to mode, pre-gated on viable CD4$^+$ cells. *Right:* MFI of FoxP3 in CD4$^+$ cells. Significance was calculated using a 2-tailed Student's t-test (*p ≤ 0.05, **p < 0.01, ***p < 0.001, ****p < 0.0001, ns = not significant).

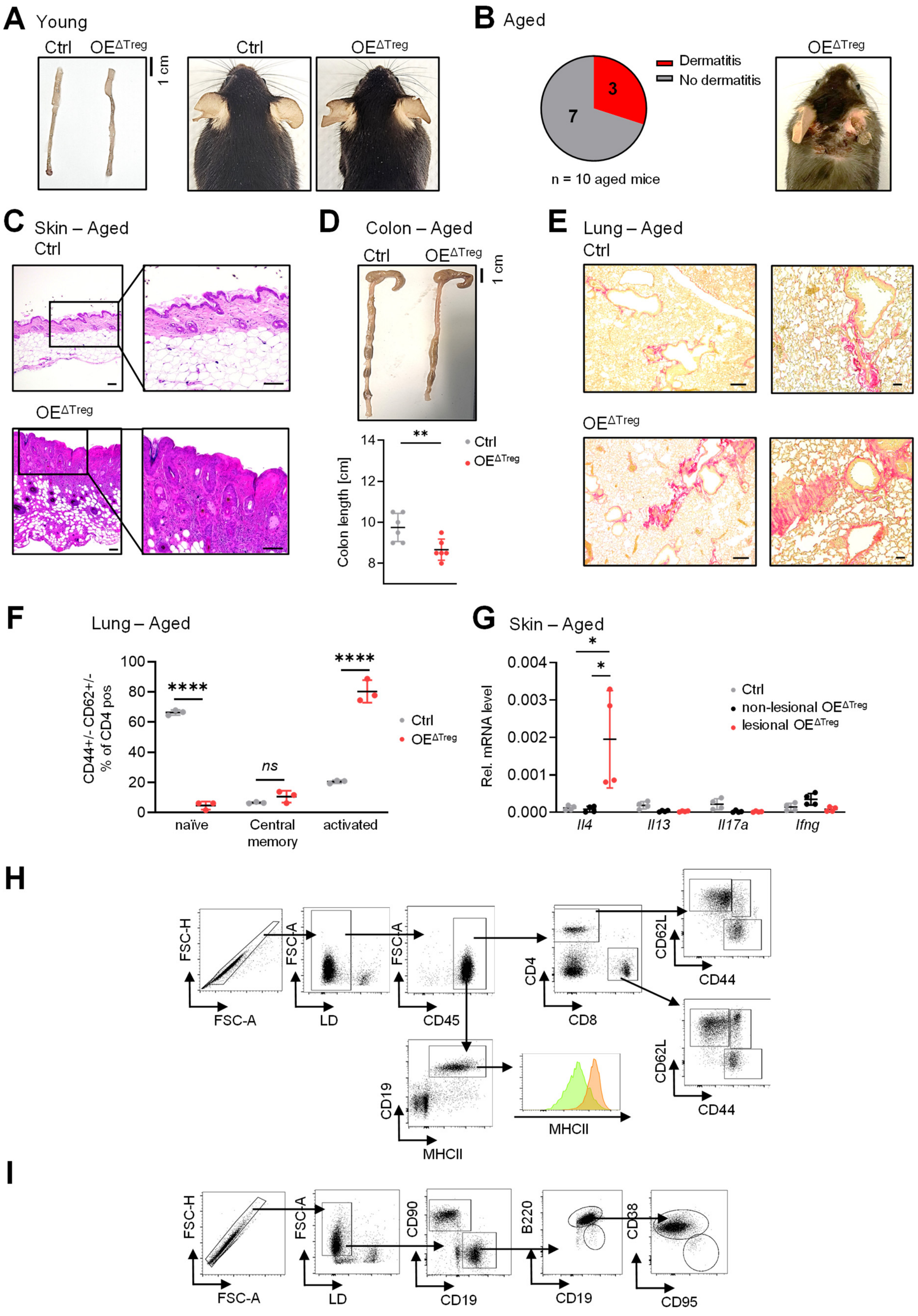


**Supplementary Figure S3. Additional data on Th2-associated disease pattern in OE$^{\Delta Treg}$ mice. A.** Images of the colon (left) and skin (right) of young (11 - 16 weeks old) Ctrl and OE$^{\Delta Treg}$ mice. **B.** Prevalence of dermatitis in aged mice (over 30 weeks) and a representative image of dermatitis in an aged OE$^{\Delta Treg}$ mouse. **C.** H&E staining of the skin from the neck of aged mice. Notably, the OE$^{\Delta Treg}$ mouse displayed overt dermatitis. Scale: 100 µm. **D.** Picture of the colon from aged Ctrl and $OE^{\Delta Treg}$ mice, and quantification of colon length. N = 6 mice per group ± SEM. **E.** Sirius Red staining of the lung from aged mice. Scale: 200 µm. **F.** Flow cytometry analysis of effector T cells in the lung of aged mice. Cells were pre-gated on viable, $CD3^+$, $CD4^+$ cells. Naïve T-cells = $CD62L^+$ $CD44^-$, central memory T-cells = $CD62L^+$ $CD44^+$, activated T cells = $CD62L^-$ $CD44^+$. N = 3 mice per group ± SEM. **G.** Gene expression in the skin of aged mice, normalized to *Actb*. N = 4 mice per group ± SEM. **H.** Gating strategy to analyze memory T cells/B cells: Following exclusion of doublets and dead cells, $CD45^+$ leukocytes were selected. Afterwards, T helper cells were characterized as $CD4^+$; cytotoxic T cells as $CD8^+$ cells. Based on CD62L and CD44 expression, naïve cells were identified as $CD62L^+$/$CD44^-$, central memory cells as $CD62L^+$ /$CD44^+$, and effector memory cells as $CD62L^-$/$CD44^+$ cells. B cells were identified based on the expression of CD19 and MHCII within the population of $CD45^+$cells. **I.** Gating strategy to analyze GC B cells: Following exclusion of doublets and dead cells, B cells ($CD19^+$) were discriminated from T cells ($CD90^+$). Within the B cell population, B1 cells ($B220^-$) were discriminated from B2 cells ($B220^+$). Within the B2 population, GC B cells were characterized as $CD38^-$ and $CD95^+$. Significance was calculated using a 2-tailed Student's t-test (*$p \leq 0.05$, **$p < 0.01$, ****$p < 0.0001$, *ns* = not significant).

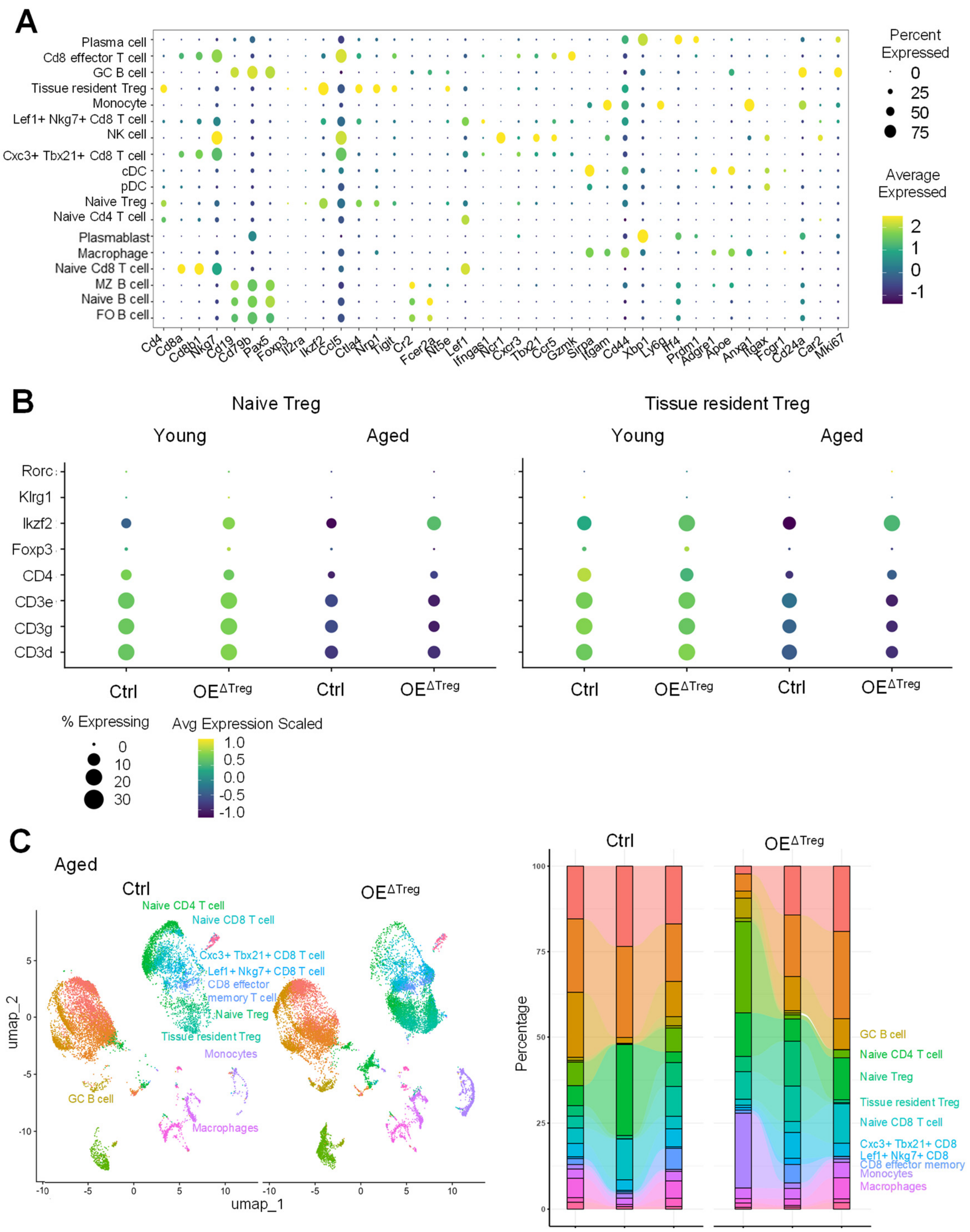


**Supplementary Figure S4. Additional single-cell sequencing data of spleens from young and aged Ctrl and OE$^{\Delta Treg}$ mice. A + B.** Marker genes used for the annotation of the identified cell clusters in the data set. **A.** General cluster annotation. **B.** General markers for the Treg clusters across all experimental groups. **C.** UMAP plot of the identified cell types in the spleen of aged ctrl and OE$^{\Delta Treg}$ mice. **D.** Relative numbers and distribution of the identified cell types in all sequenced aged mice (3 Ctrl and 3 OE$^{\Delta Treg}$). Shown is the relative percentage of the respective cell type calculated on the total number of sequenced cells.

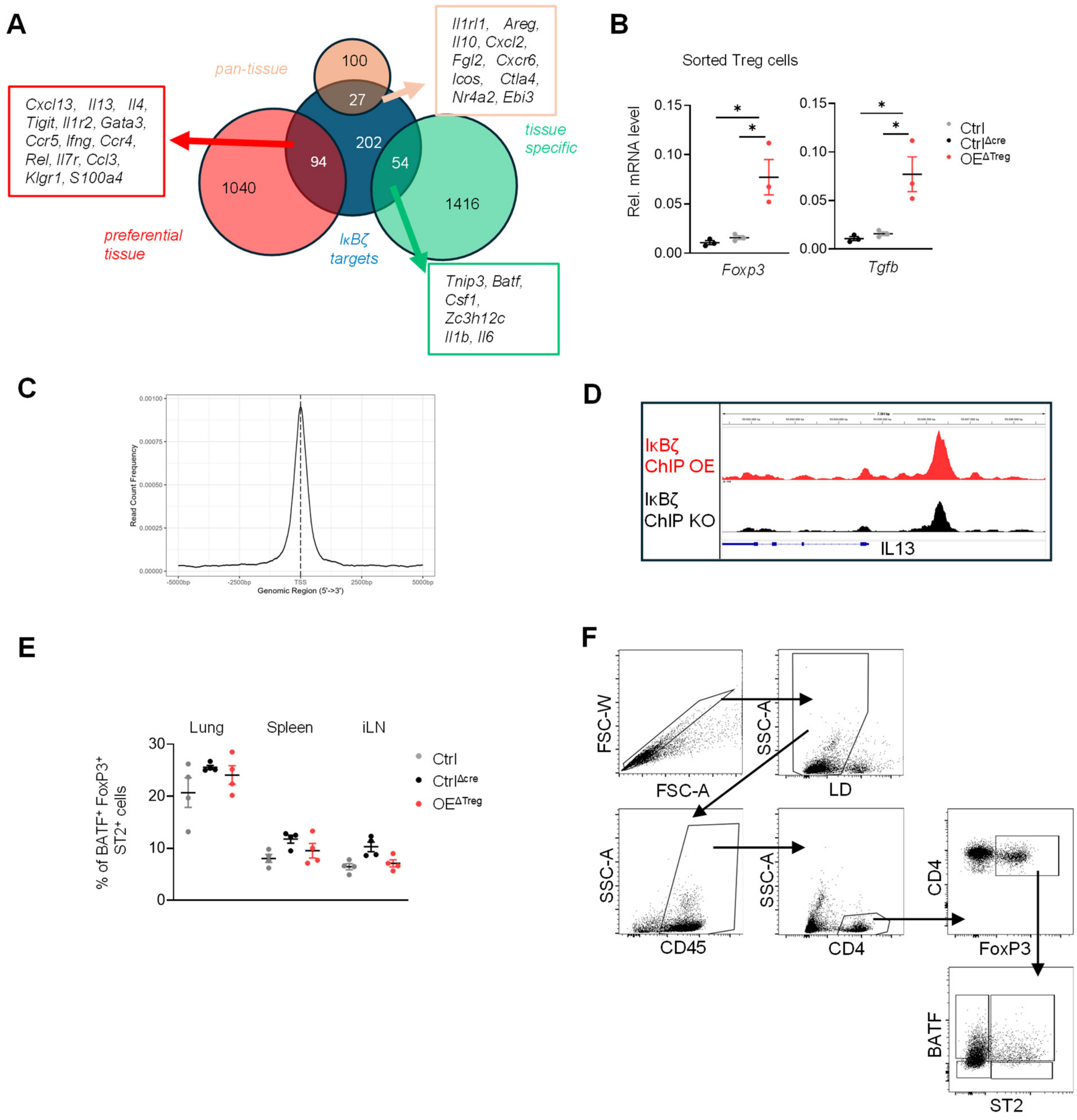


**Supplementary Figure S5. Supporting data on the modulation of BATF by IκBζ in Treg cells. A.** Treg marker genes for pan-tissue, tissue-specific, and preferential tissue-resident Treg cells, retrieved from a previously published data set [28]. **B.** Relative gene expression of *Foxp3* and *Tgfb1* in Ctrl, Ctrl$^{\Delta cre}$, and OE$^{\Delta Treg}$ sorted Treg cells. Relative mRNA levels were normalized to *Actb*. N = 3 biological replicates ± SD. **C.** Genomic distribution of IκBζ-bound regions in Treg cells, relative to the transcription start site. **D.** Screenshot of the ChIP sequencing results at the IL13 promoter region. **E.** BATF protein levels determined by intracellular FACS-staining in viable FoxP3$^{+}$ and ST2$^{+}$ cells from the spleen, inguinal lymph nodes, and the lung of young Ctrl, Ctrl$^{\Delta cre}$, and *Nfkbiz* OE$^{\Delta Treg}$ mice. N = 4 mice per group ± SEM. **F.** Gating strategy to analyze ST2$^{+}$ BATF$^{+}$ Treg cells. Following exclusion of doublets and dead cells, CD45$^{+}$ cells were selected. Within this population, CD4$^{+}$ cells were gated, and Treg cells were defined as FoxP3$^{+}$. Afterwards, BATF and ST2 expression was analyzed in CD4$^{+}$ FoxP3$^{+}$ cells. Significance was calculated using a 2-tailed Student's t-test (*$p \leq 0.05$).

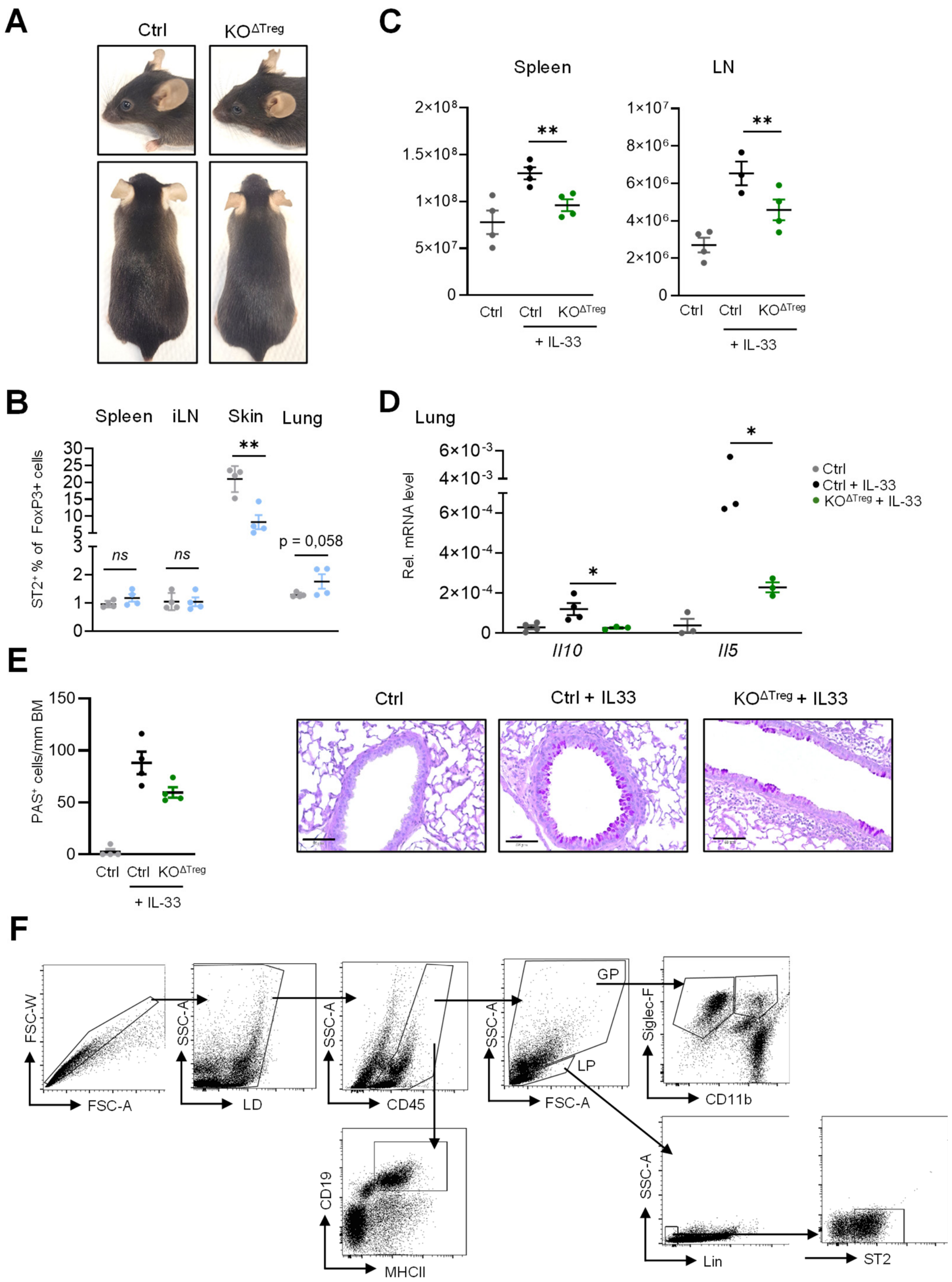


**Supplementary Figure S6. Extended characterization of *Nfkbiz* KO mice (KO$^{\Delta Treg}$) with and without IL-33 treatment. A.** Representative images of control and KO$^{\Delta Treg}$ mice. **B.** Relative frequencies of ST2$^{-}$ FoxP3$^{+}$ Treg cells in the spleen, inguinal lymph node (iLN), skin, and lung of control and KO$^{\Delta Treg}$ mice. Data show the percentage of ST2$^{+}$ cells among all FoxP3$^{+}$ Treg cells. N = 4 mice per group ± SEM. **C.** Relative number of CD45$^{+}$ cells in the spleen and iLN of control and KO$^{\Delta Treg}$ mice. N = 4 mice per group ± SEM. **D.** Gene expression analysis of skin and lung from control and KO$^{\Delta Treg}$ mice, untreated and treated with IL-33. Expression levels were normalized to *Actb*. **E.** Detection of mucus-producing goblet cells in the lung by PAS staining. *Left:* Quantification of PAS$^{+}$ cells. *Right:* Representative images for PAS staining. N = 3 - 4 mice per group ± SEM. **F.** Gating Strategy Immune cells: Within the population of doublets excluded, CD45$^{+}$ living cells. B cells were characterized as CD19$^{+}$ MHCII$^{+}$ cells. Based on characteristics in FSC and SSC, a granulocyte (GP) and lymphocyte pre-gate (LP) was used to identify macrophages, eosinophils, and ILC2. Within the granulocyte pre-gate, macrophages were identified as CD11b-low/Siglec-F-high and eosinophils as Siglec-F$^{+}$ and CD11b-high cells. ILC2 cells were identified as lineage-negative, ST2$^{+}$ cells in the lymphocyte pre-gate. Significance was calculated using a 2-tailed Student's t-test (*$p \leq 0.05$, **$p < 0.01$, ****$p < 0.0001$, ns = not significant).

### Supplementary Table S1.

Flow cytometry reagents.

| REAGENT | SOURCE | IDENTIFIER | Application |
|---|---|---|---|
| **Antibodies** | | | |
| BATF-Alexa488 | Cell Signaling | Cat. 90212 | Characterization of T cell/Treg |
| CD4-PE | BioLegend | Cat. 100408 | Characterization of T cell/Treg |
| IκBζ-PercP-efluor 710 | Invitrogen | Cat. 46-6801-82 | Characterization of T cell/Treg |
| CD45-PercP | BioLegend | Cat. 103130 | Characterization of T cell/Treg |
| FoxP3-APC | ThermoFisher | Cat. 17-5773-82 | Characterization of T cell/Treg |
| ST2-BV421 | BD Bioscience | Cat. 566310 | Characterization of T cell/Treg |
| CD45-BV510 | BioLegend | Cat. 103138 | Characterization of T cell/Treg |
| CD44-FITC | BioLegend | Cat. 103006 | Characterization of T cell/Treg |
| CD25-Pe-Cy7 | BD Bioscience | Cat. 552880 | Characterization of T cell/Treg |
| CD8-APC | BioLegend | Cat. 100712 | Characterization of T cell/Treg |
| CD62L-BV421 | BioLegend | Cat. 104436 | Characterization of T cell/Treg |
| CD45-BV510 | BioLegend | Cat. 157219 | Characterization of T cell/Treg |
| F4/80-FITC | BioLegend | Cat. 123108 | Immune cell characterization |
| CD19-PE | BioLegend | Cat. 115508 | Immune cell characterization |
| CD19-PE-Cy7 | BD Bioscience | Cat. 552854 | Immune cell characterization |
| B220-PerCP | BD Bioscience | Cat. 553093 | Immune cell characterization |
| B220-BV786 | BD Bioscience | Cat. 563894 | Immune cell characterization |
| IgM-FITC | BD Bioscience | Cat. 553437 | Immune cell characterization |
| CD11c-PE | BioLegend | Cat. 117307 | Immune cell characterization |
| SiglecF-PercP-Cy5.5 | BioLegend | Cat. 155525 | Immune cell characterization |
| ST2-APC | BioLegend | Cat. 145305 | Immune cell characterization |
| hematopoietic lineage cocktail-eFluor450 | Invitrogen | Cat. 88-7772-72 | Immune cell characterization |
| MHCII-Pacific Blue | BioLegend | Cat. 107620 | Immune cell characterization |
| CD95-PE | ThermoFisher | Cat. 12-5892 | Characterization of B cells |
| CD38-APC | ThermoFisher | Cat. 17-0381 | Characterization of B cells |
| CD19-APC-R700 | BD Horizon™ | Cat. 565473 | Characterization of B cells |

| | | | |
|---|---|---|---|
| B200-BV786 | BD Horizon™ | Cat. 563894 | Characterization of B cells |
| CD95-BV650 | BD OptiBuild™ | Cat. 740507 | Characterization of B cells |
| CD38-BUV563 | BD OptiBuild™ | Cat. 74127 | Characterization of B cells |
| CD90-BB790 | BD Bioscience | Custom Design | Characterization of B cells |
| CD90-B810 | BD Bioscience | Custom Design | Characterization of B cells |
| CD8-PE-Cy7 | BioLegend | Cat. 100722 | Isolation of Treg cells |
| CD25-PE | BioLegend | Cat. 102008 | Isolation of Treg cells |
| PD1-BV605 | BioLegend | Cat. 125220 | Isolation of Treg cells |
| TCRß-BV785 | BioLegend | Cat. 109249 | Isolation of Treg cells |
| CD11b-APC | BioLegend | Cat. 101212 | Isolation of Treg cells |
| CD11c-APC | BioLegend | Cat. 117310 | Isolation of Treg cells |
| CD19-APC | BioLegend | Cat. 115512 | Isolation of Treg cells |
| MHCII-APC | BioLegend | Cat. 107614 | Isolation of Treg cells |
| CD206-APC | BioLegend | Cat. 141708 | Isolation of Treg cells |
| NK1.1-APC | BioLegend | Cat. 108710 | Isolation of Treg cells |
| CD45-BUV737 | BD Bioscience | Cat. 752414 | Isolation of Treg cells |
| IL-33R-BV711 | BD Bioscience | Cat. 745549 | Isolation of Treg cells |
| KLRG1-BB700 | BD Bioscience | Cat. 742199 | Isolation of Treg cells |
| CD4-BV421 | BD Bioscience | Cat. 566644 | Isolation of Treg cells |
| eFluor780 fixable viability dye | ThermoFisher | Cat. 65-0865-14 | Viability dye |
| AmCyan fixable viability dye | ThermoFisher | Cat. 65-0866 | Viability dye |
| R780 fixable viability dye | ThermoFisher | Cat. 65-0865 | Viability dye |
| Zombie NIR | BD Bioscience | Cat. 423106 | Viability dye |

### Supplementary Table S2

Gene expression primers.

| Primer | Forward primer (5´ - 3`) | Reverse Primer (5´ - 3`) |
|---|---|---|
| *Nfkbiz* | AACTCGCCAAGAGACCAGTG | AGAGCCACTGACTTGGAACG |
| *Ifng* | GACAATCAGGCCATCAGCAAC | CATTGAATGCTTGGCGCTGG |
| *Il4* | CATCGGCATTTTGAACGAG | CGAGCTCACTCTCTGTGGTG |
| *Il10* | GCATTTGAATTCCCTGGGTGAG | CATGGCCTTGTAGACACCTTGG |
| *Il13* | AACATCACACAAGACCAGACTCC | CCAGGTCCACACTCCATACC |
| *Il17a* | GCCCTCAGACTACCTCAACC | TTCCCTCCGCATTGACACAG |
| *Batf* | TAGAACCATGCCTCACAGCTC | TGAAGGGTGTCGGCTTTCTG |
| *Foxp3* | GGCCCTTCTCCAGGACAGA | GCTGATCATGGCTGGGTTGT |
| *Tgfb* | CCCGAAGCGGACTACTATGC | CATAGATGGCGTTGTTGCGG |
| *Actb* | AGGAGTACGATGAGTCCGGC | GGTGTAAAACGCAGCTCAGTA |
| *Areg* | GCATGCCATTGCCTAGCTG | ACCATCCGAAAGCTCCACTTC |
| *Il5* | ACATTGACCGCCAAAAAGAG | ATCCAGGAACTGCCTCGTC |

### Supplementary Table S3

ChIP primers.

| Primer | Forward primer (5´ - 3`) | Reverse Primer (5´ - 3`) |
|---|---|---|
| *Il10* | GCCAGTTAGAAAGCCACCAC | TGTGCTCATAGGCTGTCTGG |
| *Batf* | ACTTGCTAATTCTGGCCGGG | GGCAGTTTGAAAGGCAGAGC |
| *Il4* | GGAGGTTCTGGGCTAGGTTG | ACACGGTGCAAAGAGAGACC |
| *Il13* | GTGCCTTGCCTAGCCAAATG | TTCTGCTTGTCTTGAGGGGC |
| *Mb* | GATGACCCGTGTTTCGACTC | GTAGGGAAGTGTGCGTGCTC |
